# Impact of gravitational forces on Red Blood Cell dynamics in biofluid suspension

**DOI:** 10.1101/2024.04.22.590537

**Authors:** Anirudh Murali, Ram Rup Sarkar

**Affiliations:** Chemical Engineering and Process Development, CSIR - National Chemical Laboratory, Pune, India; Academy of Scientific and Innovative Research (AcSIR), Ghaziabad 201002, India

**Keywords:** Altered gravity, Red Blood Cell, fluid flow, Dissipative Particle dynamics, computational modelling

## Abstract

The growing interest in space exploration and human spaceflight has highlighted the critical challenges posed by microgravity on human physiology. One such challenge is the behavior of immune and red blood cells, which primarily rely on hematic or lymphatic streams (biofluids) for transport. This study aims to quantitatively address the puzzle of how red blood cells are influenced by gravity when they are suspended in bio-fluid. The Dissipative Particle Dynamics (DPD) approach was used to model blood and the cell by applying gravity as an external force along the vertical axis and varied from 0g to 2g during parameter sweeps. Key metrics, including Elongation and Deformation indices, pitch angle, and normalized center of mass, were utilized to assess cellular behavior. Results revealed that gravity induces shape changes and spatial alignment in red blood cells. The Elongation Index and the normalized center of mass declined linearly with the applied gravity. Correlation analysis showed a strong correlation between applied gravity and the aforementioned variables. Additionally, forces acting on the cell, such as drag, shear stress, and solid forces, diminished as gravitational force increased. Further analysis indicates that increasing gravity affected the cell’s velocity, resulting in prolonged proximity to vessel walls and intensified viscous interactions with surrounding fluid particles, thereby triggering morphological changes. This study provides crucial insights into the biophysical effects of gravity on the red blood cell and presents a significant step toward understanding cellular dynamics under altered gravitational conditions.

## 1. Introduction

In recent years, there has been a significant surge in the intrigue around space and human spaceflight. This is evident through the establishment of the Artemis Accord, which signifies humanity’s intention to dispatch personnel to both the Moon and Mars during the timeframe of the 2030s (Creech et al. 2022), necessitating an extended stay in hostile environments. However, previous studies have reported various physiological changes in astronauts who have spent extended periods in the International Space Station (ISS). These changes include heightened allergic reactions, decreased counts of T- and B-cells, impaired morphology and activation of immune cells (Crucian et al. 2008), reduced activity of macrophages (Kaur et al. 2005; Moreno-Villanueva et al. 2018), and inconsistent release of interleukins and chemokines (Hauschild et al. 2014; Moreno-Villanueva et al. 2018; Thiel et al. 2017). Given the potential health risks associated with space exploration, it is essential to conduct a thorough investigation into the influence of gravity on the immune and associated systems to ensure the well-being of humans in future space missions.

The effects of microgravity (10^−6^g; g = 9.81 m/s^2^) on the human body have been thoroughly studied, with body fluids recognized as a significant source of concern. Prisk (Prisk 2011) states, body fluids collect in the thorax before redistribution and eventual balance. While immune cells can be found as resident cells in tissues, their primary mechanism of movement to their site of action is via hematic or lymphatic streams (Hampton and Chtanova 2019). Several studies have related the absence of convection to the lack of driving forces in microgravity (except capillary force). The lack of convection reduces shear pressures (Nickerson et al. 2003), cell sedimentation (Wilson et al. 2002), nutrient delivery, and waste elimination (Guéguinou et al. 2009), resulting in decreased gene expression and subsequent physiological consequences (Crawford-Young 2006; Nickerson et al. 2003). Prior studies have suggested that these factors contribute to the immune system’s impairment and, as a result, a decrease in immunological activity. Therefore, understanding the association between cellular mechanics and mechanotransduction in microgravity is critical to solving the immune dysfunction puzzle. Microgravity research has primarily focused on experimental procedures, and studies have been done from a mechanics perspective on vesicles and RBCs (Callens et al. 2008; Mader et al. 2006; Podgorski et al. 2011) under microgravity. Similarly, some computational studies have been done on non-adherent cells such as stem cells (Ghaemi et al. 2021) and bone/osteocytes (Liu et al. 2022; Liu et al. 2020; Wang et al. 2022; Zhao et al. 2020); their underlying biology has not been investigated.

The mechanism through which cells perceive gravity remains a puzzle. While certain structures like the Otolith in the ears are confirmed to play a role, the exact process by which non-adherent cells sense gravitational forces, especially in conjunction with shear forces in fluid suspension, remains uncertain. Questions linger regarding whether these cells uniformly sense gravity across their surfaces or if specialized structures on their surfaces are involved. If direct sensing occurs, is there a specific point of contact inside the cell to transmit this force? Alternatively, if a protein on the cell surface mediates this process, what kind of structural change enables it to relay signals internally? Additionally, the unclear understanding of how cells respond to mechanotransduction under conditions of microgravity and shear further complicates the picture. These questions are an unresolved topic on the complex interplay between physics (microgravity, forces, and cellular morphology), biology (gene regulation), and chemistry (small molecule production), underlined in a recent review (Murali and Sarkar 2023).

In this study, we developed a numerical model based on Dissipative Particle Dynamics (DPD) to investigate the behavior of red blood cells (RBCs) under various gravitational conditions. Building upon our previous work, which described RBC behavior under varying gravity (Murali and Sarkar 2024), this model introduces an improvised framework that couples fluid movement with solid structures through DPD-based force equations and incorporates improved boundary conditions to account for wall fluctuations. This approach addresses broader questions regarding how cells undergo shape changes and respond to shifts in gravity. The primary objective of this study was to investigate shape alterations induced by external forces and to identify the factors driving these changes using metrics such as the Elongation Index, Deformation Index, pitch angle, and normalized center of mass. We employed a gyration tensor-based approach to estimate and elucidate the shifts associated with these metrics to achieve this. Additionally, we examined the effects of mechanical forces, including drag, shear stress, and other solid forces, on RBC behavior in response to gravitational variations. The results imply that forces are influenced by cell migration speed, interaction with surrounding fluids, and the magnitude of applied gravity. Our findings collectively elucidate how cells respond to varying gravitational forces, detailing their movement, interactions with vessel walls, and the often challenging-to-measure forces they experience during these processes in experiments.

### 2. Material and methods

This section provides a concise overview of the dissipative particle dynamics (DPD) formulation with a two-dimensional discrete cell model of red blood cells (RBC).

### 2.1. Bio-Fluid model

DPD is a mesoscopic scale simulation technique initially proposed by Hoogerbrugge and Koelman (Hoogerbrugge and Koelman 1992) that utilizes particles as its fundamental representation units. The forces exerted on the particle are determined by evaluating the pairwise distances between particles within a specified cut-off radius. The DPD method computes three internal forces for each particle while exerting an external force on each particle. Newton’s second law provides a framework for understanding the temporal progression of forces,

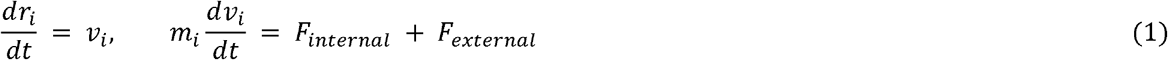

Where *r*_*i*_ and *v*_*i*_ are the position and velocity vectors of particle i, and the unit of mass is taken to be the mass of a particle. F_Internal_ is the internal forces generated due to pairwise interactions between particles. Internal forces can be categorized into three types: conservative force (F^C^), dissipative force (F^D^), and random force (F^R^). Conservative force is a linear soft repulsive potential; dissipative force is a viscous force to mimic viscous friction, and the random force accounts for the Brownian motion at the meso- and microscopic scale.

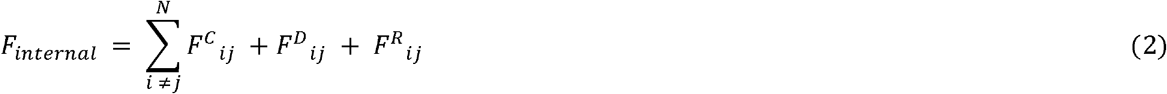

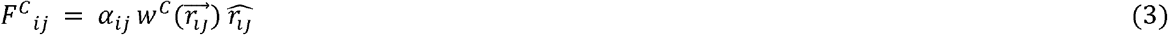

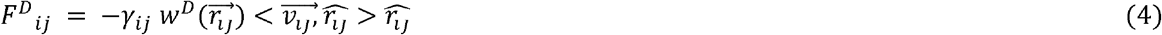

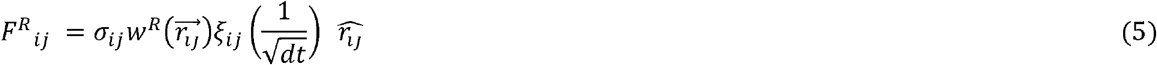

The strength of dissipative and random forces is represented by the dissipative strength parameter (*γ*_*ij*_) and the random strength parameter (*σ*_*ij*_), respectively. The soft repulsive potential (**Eq. 3**) and conservative force coefficient (*α*_*ij*_), keep interacting particles from getting too near to one another. *r_ij_* is the distance between particle i and j, 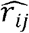 is its unit vector, and 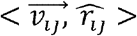 is the dot product of the unit vector and velocity difference between particle i and j. *ξ*_*ij*_ is a random number following zero mean and unit variance and has the following properties.

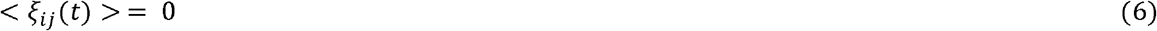

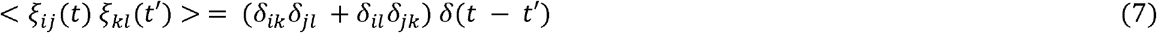

The random number applied to a pair of particles is kept constant to preserve momentum. Interparticle-dependent weight parameters *w*^*C*^, *w*^*R*^, *w*^*D*^ vanish at 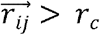.

These weight parameters are as follows

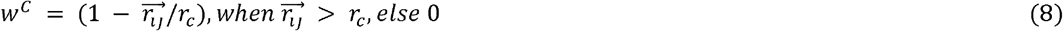

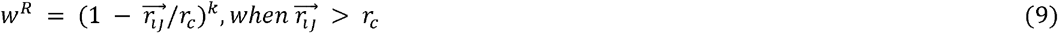

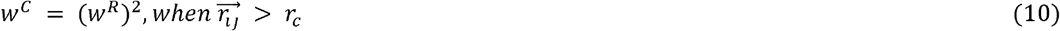

Groot and Warren (Groot and Warren 1997) found a relationship between the dissipative strength parameter, random strength parameter, and the fluctuation-dissipation theorem given by,

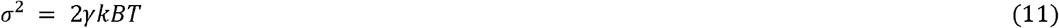

kBT stands for the system temperature. As the random force tends to “heat up,” while the dissipative force seeks to cause viscous damping in the system, doing so will guarantee the temperature stability of the system. The corresponding dissipative and random parameter in **Eq. 11** explains how the two forces work as a thermostat in this way.

#### Boundary conditions

Similar to Fan et al. (Fan et al. 2003), to implement the no-slip boundary condition, particles that penetrate the wall layer are reflected back into the simulation domain, whereby their velocity is reassigned from a normal distribution. Cartesian coordinate system is chosen as a frame of refence, with x-direction along the horizontal and the y-direction along the vertical axis (shown in **Fig. 1**). The direction of reflection is determined by the vector ‘n’, which is orthogonal to the boundary. The periodic boundary condition is enforced in the x-direction.

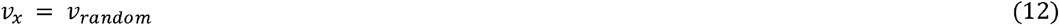

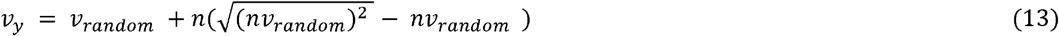

**Fig. 1:**
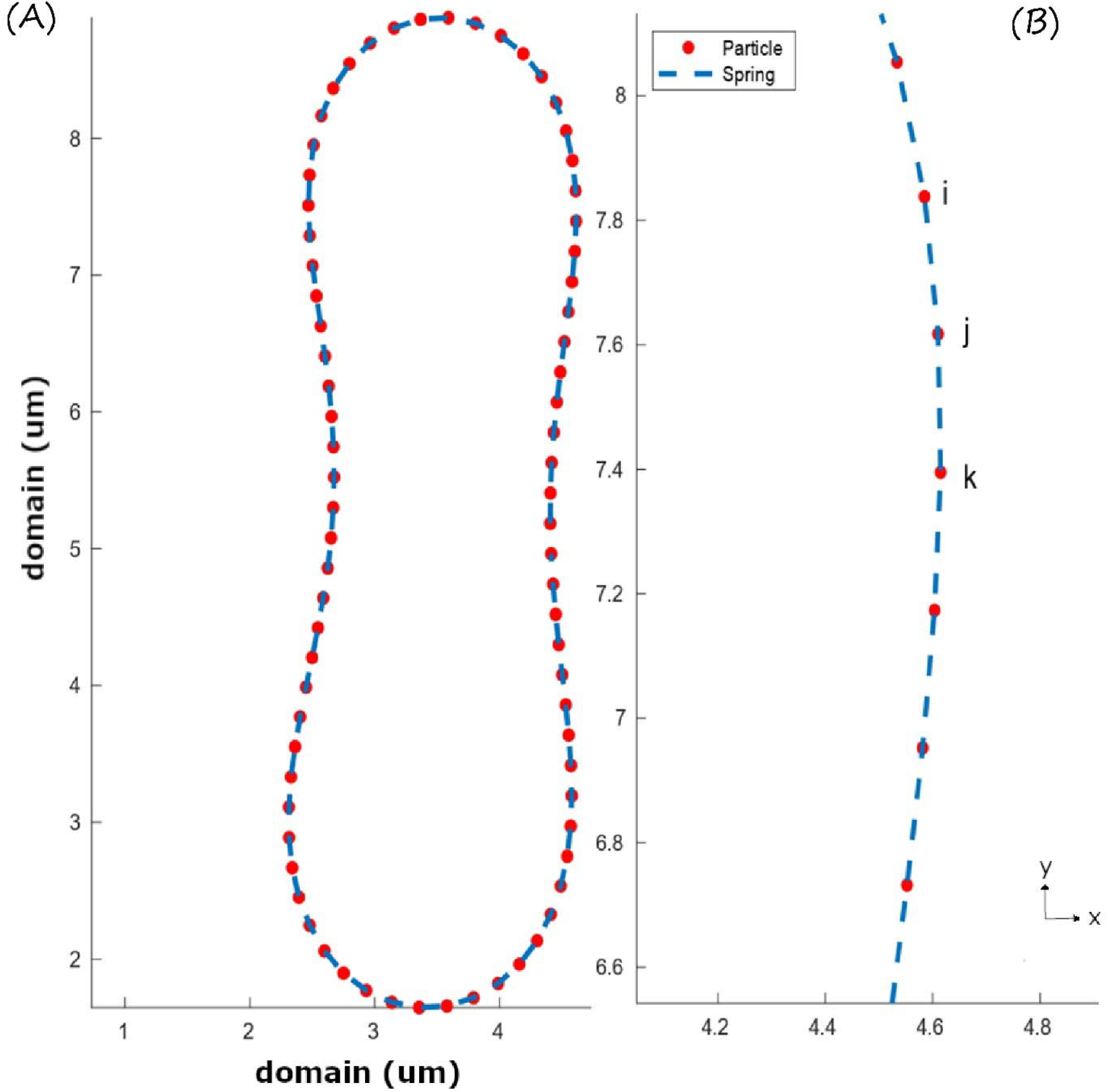
(A) Minimum energy shape of RBC when the reduced area parameter is applied. (B) Nodes i, j, and k are connected by a spring of length l_0_ at rest and the intersection of two adjacent springs, ij and jk, at the junction point j subtends angle *θ*_*ijk*_.

#### Parameter scalin

DPD simulation is carried out in reduced units. In order to scale parameters, defining the reference parameters becomes necessary. In this study, the physical reference length and cut-off radius (r_c_) are 1×10^−6^ m; in DPD terms, r_c_ equals 1. Particle density (ρ_DPD_), reference temperature (T), reference time (t_ref_), reference velocity (v_ref_), and Young’s modulus of RBC are additional reference parameters required to execute the DPD simulation. The **Supplementary Information, Section 1, Parameter derivation**, provides the derivation of simulation parameters.

### 2.2 Cell model

The mean diameter of a human RBC ranges from 6.10 to 8 micrometers. When fully developed, it assumes a biconcave morphology and exhibits an absence of a cellular nucleus. The structure of RBCs consists of phospholipid bilayers and the cytoplasm. Of which the cytoplasm can be described as a viscous Newtonian liquid. The cytosol preserves the internal volume and uniformly disperses external pressures exerted on its surface. The RBC membrane consists of a spectrin network interconnected with the cell wall. The structural networks, which govern the general deformation behavior of RBCs, constitute most of the load-bearing framework of RBCs. The overall shape is determined by various factors, including the cell membrane’s elasticity, surface area, and the volume enclosed within the cell. Thus, RBCs possess a certain degree of flexibility, like a partially inflated capsule, and that shape can be used to model it.

In this study, RBC is modeled as a ring of 76 finite-sized DPD particles interconnected by non-linear Worm Like Chain (WLC) springs. Multiple studies have employed different numbers of particles (ranging from 10 to 88) to simulate RBC (Hoque et al. 2018; Pan et al. 2010; Polwaththe Gallage et al. 2014; Tsubota et al. 2006). However, the present formulation is similar to Pan et al. (Pan et al. 2010); the spring is modeled as a worm-like chain (WLC), and an area penalty function is applied to conserve the RBC shape and area (Tsubota et al. 2006). RBC initially starts as a circle with a radius of 3.05r_c_; applying the area penalty function forces the RBC to settle to its minimum energy shape. The energy functions are presented below:

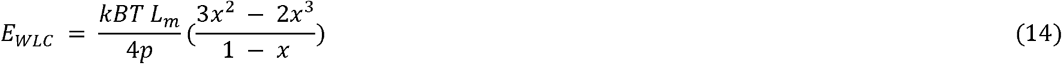

E_WLC_ represents elastic energy stored by a spring described by a non-linear relation, where kBT is the system temperature, L_m_ is the maximum permissible length of the spring, p is the persistence length of the flexible spring, and x is the ratio L_j_/L_m_, where L_j_ is the spring’s current length. Similarly, to account for the membrane’s bending and compressional stiffness. A three-point interaction potential (E_bend_) is applied where k_b_ is the RBC membrane bending constant, and *θ*_ijk_ is the instantaneous angle between the two consecutive springs with the j^th^ particle as the center.

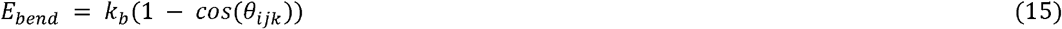

To maintain shape and conserve the internal, we apply the area penalty function as a potential,

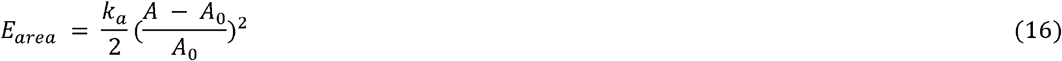

Where k_a_ is the area coefficient, A is the instantaneous area, and A_0_ is the intended area of RBC. The reduced area parameter is introduced to define the intended enclosed area, and the relationship is given by 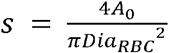. The total energy of the system and the forces is given by

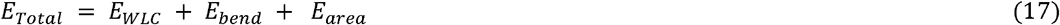

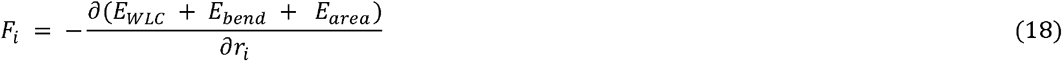

The values of the parameters described above are shown in **Table 1**.

**Table 1:**
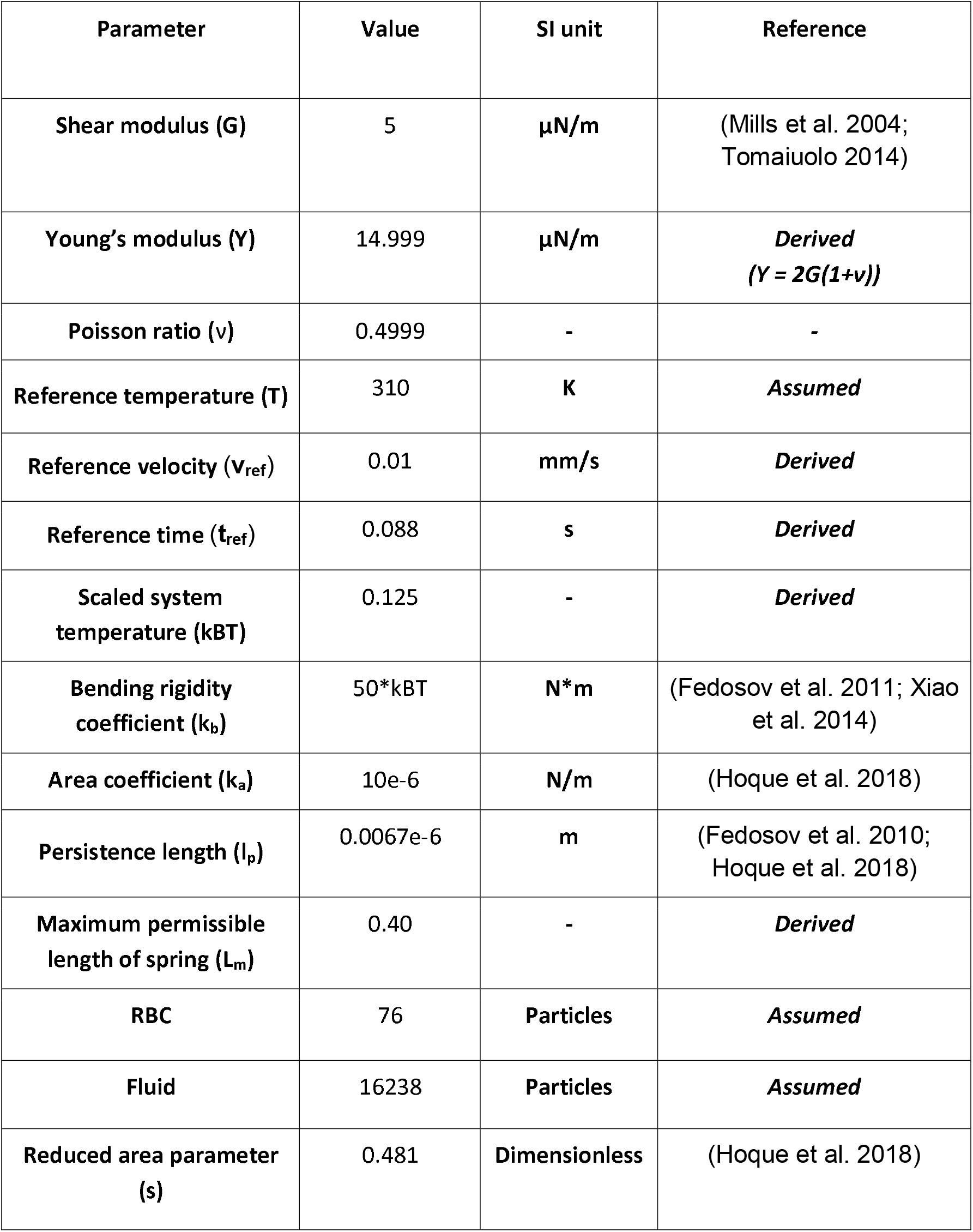
List of all physical parameters used in the DPD simulation.

#### Computational details and Benchmarking

The parametric study of fluid flow and structure interaction is conducted at a physiological temperature of 310 K, which is equivalent to 0.125 in DPD units, corresponding to the average temperature of the human body, and a cutoff radius of 1r_c_ (DPD units), representing a microscopic dimension. The fluid particles were initially populated and arranged in a Face-Centered Cubic (FCC) unit cell configuration. Based on the dimensions of the simulation box and the specified arrangement, the total number of particles was determined to be 16238, and the corresponding number density (ρ_DPD_ = total particles/dimensions of simulation box (202.5r_c_ x 20r_c_)) is estimated to be around 4.00. Periodic boundary conditions (PBC) are applied at far end of the x-axis to allow the flowback of particles and reflection with repulsion implemented at the walls to mimic the no-slip condition. The particles are initialized with random values for velocity and acceleration. The DPD system can be classified as an NVE (Particle, Volume, and Energy) ensemble, in which the three entities are conserved and system temperature is monitored for fluctuation. The minimized RBC shape, as depicted in **Fig. 1**, is positioned at the center of the left boundary. The fluid particles contained within the RBC are regarded as the RBC cytosol. In order to induce flow, a minor external force is applied to the system in the x direction, and to mimic gravitational conditions, another eternal force is applied in the y direction representing gravity. The dissipative strength parameter (*γ*_*ij*_) is chosen to be 4.5, and the random strength parameter (*σ*_*ij*_) is calculated to be 0.795 based on the relation in **Eq. 11**. The Conservative force coefficient (*α*_*ij*_) is considered 2.34 based on the following relation 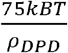 given by Groot and Warren (Groot and Warren 1997).

Simulations were performed for 25 seconds in the physical scale, with a timescale equivalent to 0.088 sec and a time step size of order O (10^−3^). An in-house Python-based code was developed to implement a two-dimensional model for blood flow, and data analysis was carried out in MATLAB 2020a. The interaction characteristics pertaining to various types of particles are presented in **Table 2**.

**Table 2:**
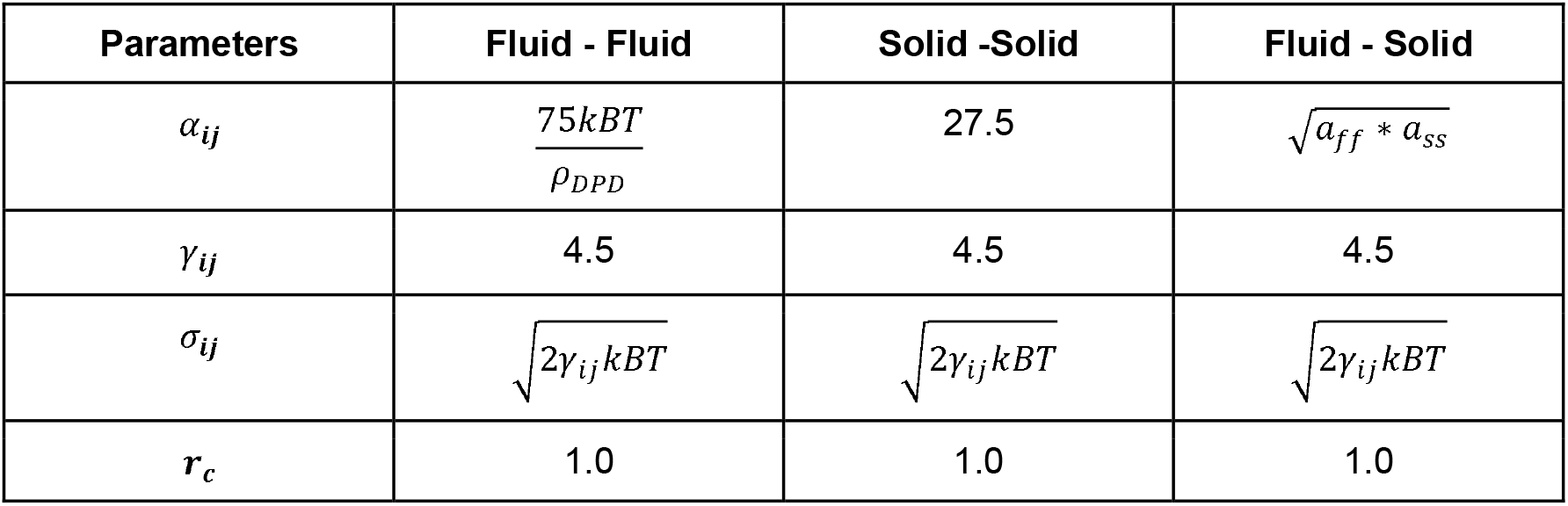
DPD interaction parameters.

### Statistical analysis

The data and subsequent statistical analyses were performed utilizing MATLAB® 2020 software. Evaluation of cell morphology and spatial alignment involved fitting normal probability distribution functions to the aggregated data using the fitdist package. Hypotheses regarding the potential differences in the median values of parameters including EI, DI, theta and YCoM between the 1g sample and both micro- and hypergravity samples were assessed via two-sided nonparametric test (Mann-Whitney U test, ranksum package), employing a dataset comprising 5112 samples from 6 simulation runs for each gravitational condition. Significance levels were designated as p < 0.05, p < 0.01 and p < 0.001 to denote statistical significance between the compared samples.

## 3 Results and discussions

### RBC mechanics

A stretch test is performed to evaluate RBC’s structural integrity and mechanical properties of RBC. In this study, we follow the protocol laid out by Dao et al. (Dao et al. 2003). RBC is affixed to a silica bead at one extremity and subjected to incremental force to induce displacement from the opposing end. This test enables the assessment of its strength and the extent of deformation that may arise due to external forces. The variations in axial and transverse diameter are measured and graphed in relation to the applied force (**Fig. 2(A)**). These measurements are then compared to the findings of Bohiniková et al., Fedosov et al., and Suresh et al. (Bohiniková et al. 2021; Fedosov et al. 2010; Suresh et al. 2005). The present simulation agrees with the experimental result and the 3D simulations. Similarly, we compare the changes in RBC shape during the optical tweezer test of Zhang and Liu (refer to Figure 3(c)) (Zhang and Liu 2008) at specific force intervals **(Fig. 2(B)**). The shape change observed in our simulation is comparable to the shape produced during the experiment. Based on the agreement of the results, parameters used during the stretch test were used in the DPD simulation.

**Fig. 2:**
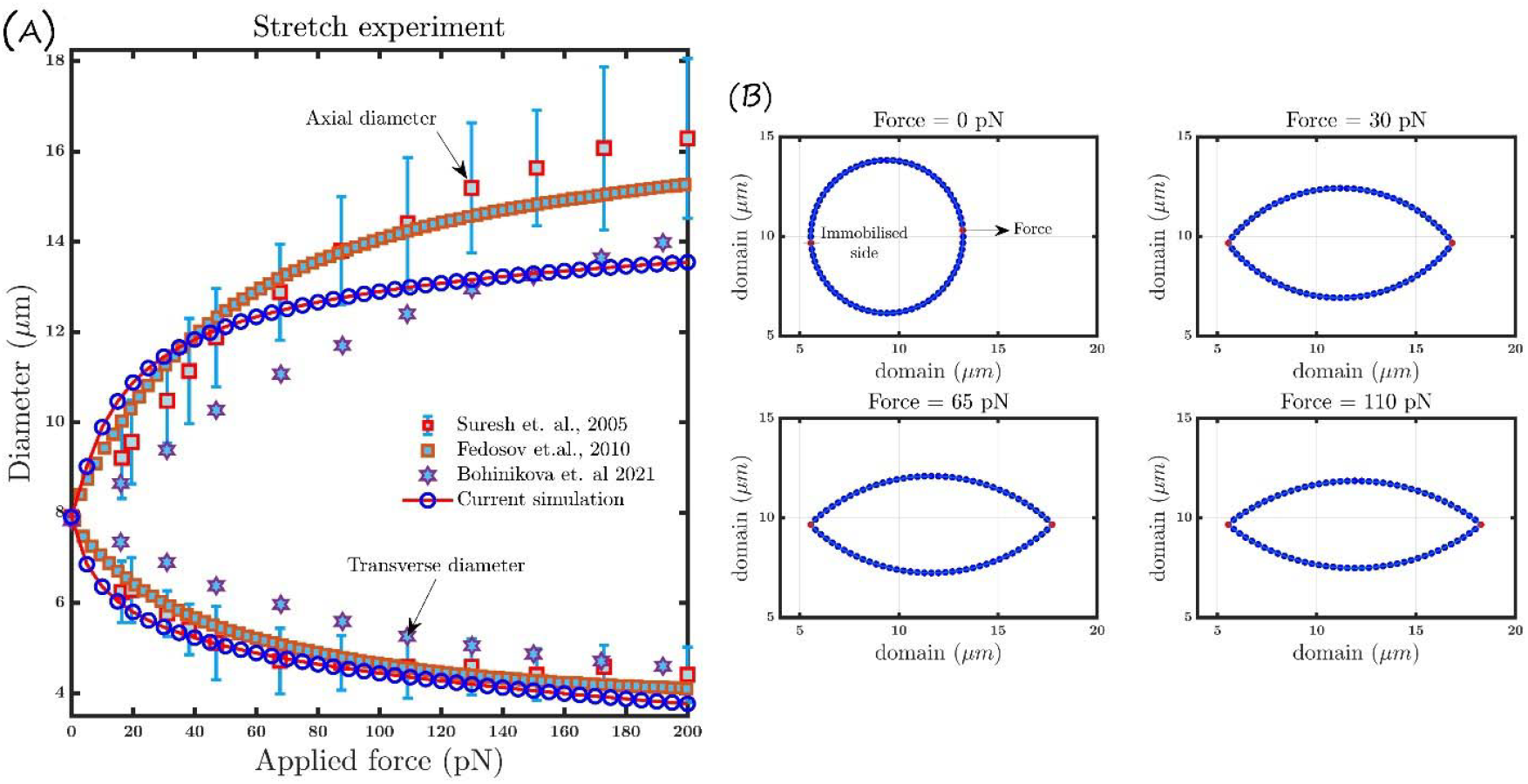
(A) Axial and transverse diameter of the RBC when deformed with external stretch force, Current (blue circle) compared to Suresh et al., (Suresh et al. 2005) experimental test, Bohiniková et al., (Bohiniková et al. 2021) and Fedosov et al. ‘s (Fedosov et al. 2010) 3D simulation results. (B) Shapes produced at specific force intervals in the optical tweezer test closely resemble Zhang and Liu’s optical tweezer test (Zhang and Liu 2008).

**Fig. 3:**
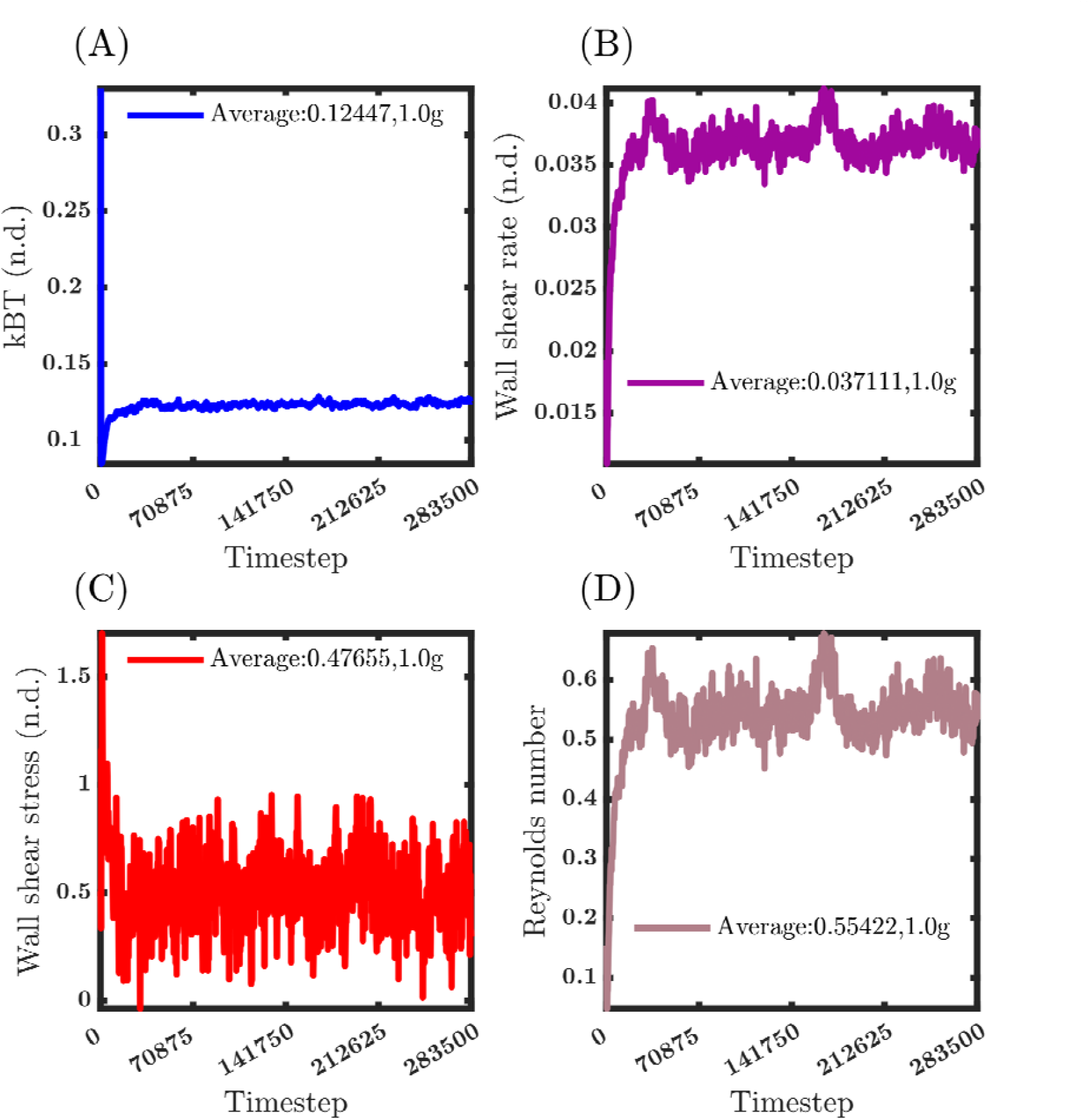
Temporal evolution of various parameters during the simulation under 1.0g condition; value averaged for the last 25% of data. (A) Temperature fluctuations (kBT) stabilize after 10,000 timesteps. (B) The wall shear rate fluctuates before stabilizing after 39,150 timesteps. (C) Wall shear stress rises at the start of the simulation and eventually reaches equilibrium. (D) Reynolds number initially increases and stabilizes after 39,150 timesteps.

### Cell dynamics and parametric sweep

To investigate the interactions between fluids and solids at the mesoscale, the cell was energy-minimized and then placed at the start of the fluid domain, which had previously reached a state of equilibrium. With the introduction of RBC into the system, a spike in temperature was observed, eventually stabilizing near the equilibrium kBT value. Once systemic equilibrium with the RBC is achieved, the cell experiences forces, such as shear and drag, from the surrounding fluid. To explore the impact of gravity on cellular behavior, gravity was introduced as an external force along the y-axis and varied from 0g (< 1g - microgravity regime) to 2g (> 1g - hypergravity regime) in an increment of 0.25g. The system was simulated over a 25-second timeframe on the physical scale. The computational domain was segmented into 400 bins in the y-direction for further analysis to calculate rheological and cell-dependent properties. Local flow parameters were determined by averaging each bin’s sampled data from the last 25% of the data. It was observed that, over time, due to the interaction with the surrounding fluid, RBCs transitioned from a biconcave shape to various configurations, such as those resembling a symmetric parachute, slipper, asymmetric parachute, or anti-parachute. The aforementioned configurations that RBC assumed were contingent upon its location within the given domain. In order to determine the transient shape change, two distinct parameters were introduced, namely the elongation index and the deformation index, derived from the Gyration tensor which are expressed as follows:

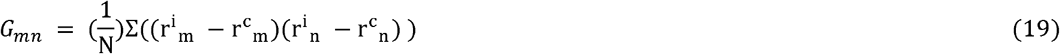

The notation *r*^*i*^_*m*_, *r*^*i*^_*m*_ denotes the current position of the solid vertex (particle) and *r*^*c*^_*m*,_ *r*^*c*^_*n*_ represents the center of mass of the solid in the x and y directions, respectively. The gyration tensor is a 2 × 2 tensor; diagonalizing the tensor gives two eigenvalues, λ _1_ and λ _2_. These eigenvalues represent the cell diameter along the x and y directions.

Assuming, *λ*_1_ > *λ*_2_, the ratio of these eigenvalues gives,

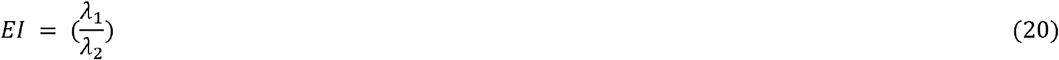

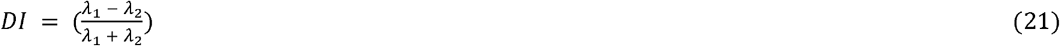

To analyze the impact of the surrounding fluid and gravity on a cell, drag, shear, and acceleration acting on a solid object were calculated based on Chen et al.’s (Chen et al. 2006) formulation.

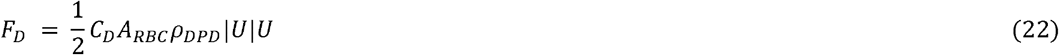

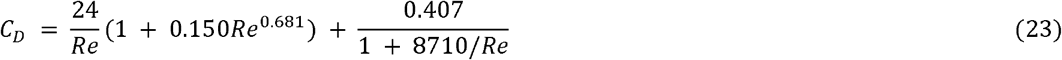

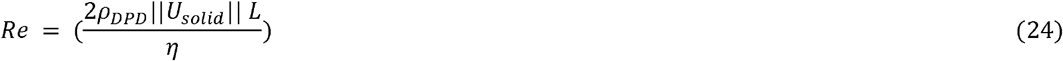

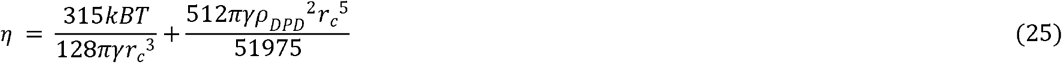

In this context, F_D_ represents the drag force, C_D_ denotes the drag coefficient, ***A_RBC_*** signifies the area enclosed by RBC, ***ρ_DPD_*** represents the system’s density, and U denotes the RBC’s velocity. Viscosity (***η***) was estimated using the methodology outlined in Fan et al. (Fan et al. 2006) to calculate the drag coefficient. Eq. 23 establishes a scaled correlation between the Reynolds number (ratio of inertial to viscous forces) and the drag coefficient; Brown and Desmond (Brown Phillip P. and Lawler Desmond F. 2003) established a correlation by analyzing sphere drag data collected over the twentieth century, valid for Reynolds numbers below 100,000. The calculated values were compiled and categorized based on temporal and parametric variables (presented in **Fig. 3** and **Supplementary Information, Section 2, Fig. S1-S8**).

Furthermore, the pitch angle and normalized the longitudinal center of mass of the cell were monitored in addition to assessing elongation and deformation indices. The phenomena of elongation and deformation indices offer insights into the alterations in shape resulting from the contact between a fluid and a solid. The former two can be utilized to assess the physical changes that impact the cell due to gravitational changes. Time-dependent variations of all the previously mentioned indices are presented in **Fig. 4**.

**Fig. 4:**
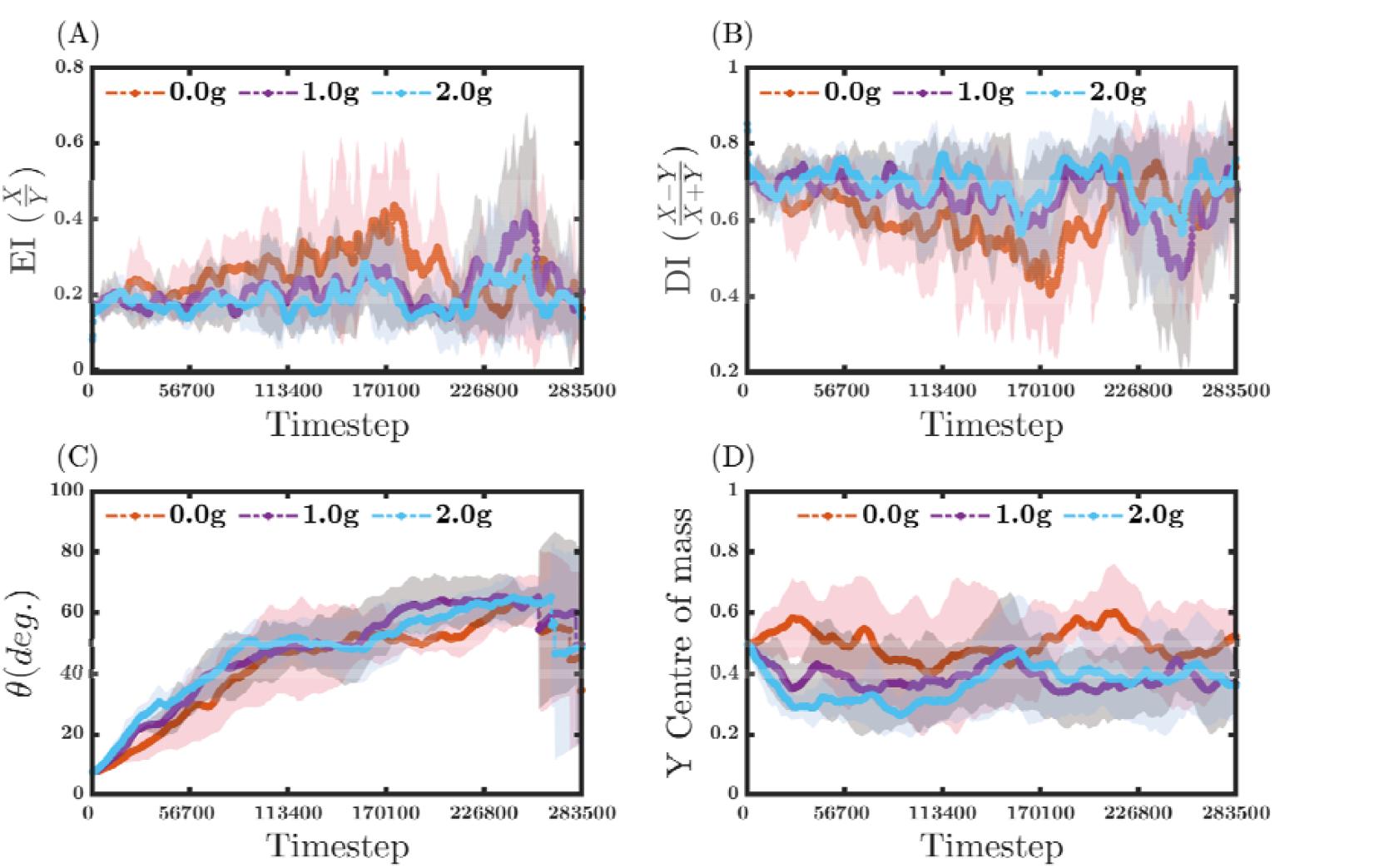
Temporal evolution of key solid parameters under varying gravitational conditions (0.0g, 1.0g, and 2.0g) presented with their standard deviation in the shaded region. (A) The elongation index (EI) for each gravitational condition indicates fluctuations but cells under microgravity show higher EI for longer period of time. (B) The deformation index (DI) for each gravitational condition indicates higher deformation to at 0g when compared to other gravity conditions (C) RBC pitch angle (θ) shows a progressive increase, with variations depending on the gravitational condition. (D) Y center of mass (YCoM) demonstrates a decline under higher gravity levels and however due to RBC’s structure it comes back to the centerline at 2g condition, whereas at 0g the cell tends to stay along the centerline of the domain.

In **Fig. 4**, it is evident that the cell subjected to higher gravity (hypergravity in comparison to 1g) exhibits subdued elongation but higher deformation, rapid shift in pitch angle, and movement toward the bottom wall of the domain. Conversely, the cell exposed to reduced gravity (microgravity) experienced the opposite effects. While it is possible to derive specific insights by examining the temporal fluctuations of these indices, however the application of the random force induces stochasticity in the system, contributing to noise in the temporal scale. Therefore, to comprehensively derive an understanding of the temporal behaviors, aggregation of data becomes necessary to identify fundamental distribution patterns. Hence, to ascertain the distribution and likelihood of these quantitative metrics, probability histograms were constructed, and theoretical distributions were fitted over the probability histogram data (**Fig. 5 and 6, Supplementary Information, Section 2, Fig. S9 and S10**). Various theoretical distributions were used to fit the constructed histograms. Among them, the normal distribution fit was able to explain few of the histograms, but majority of the histograms did not follow normal distribution. Therefore, subsequent hypothesis testing employed a non-parametric approach to avoid assumptions about the underlying distribution.

**Fig. 5:**
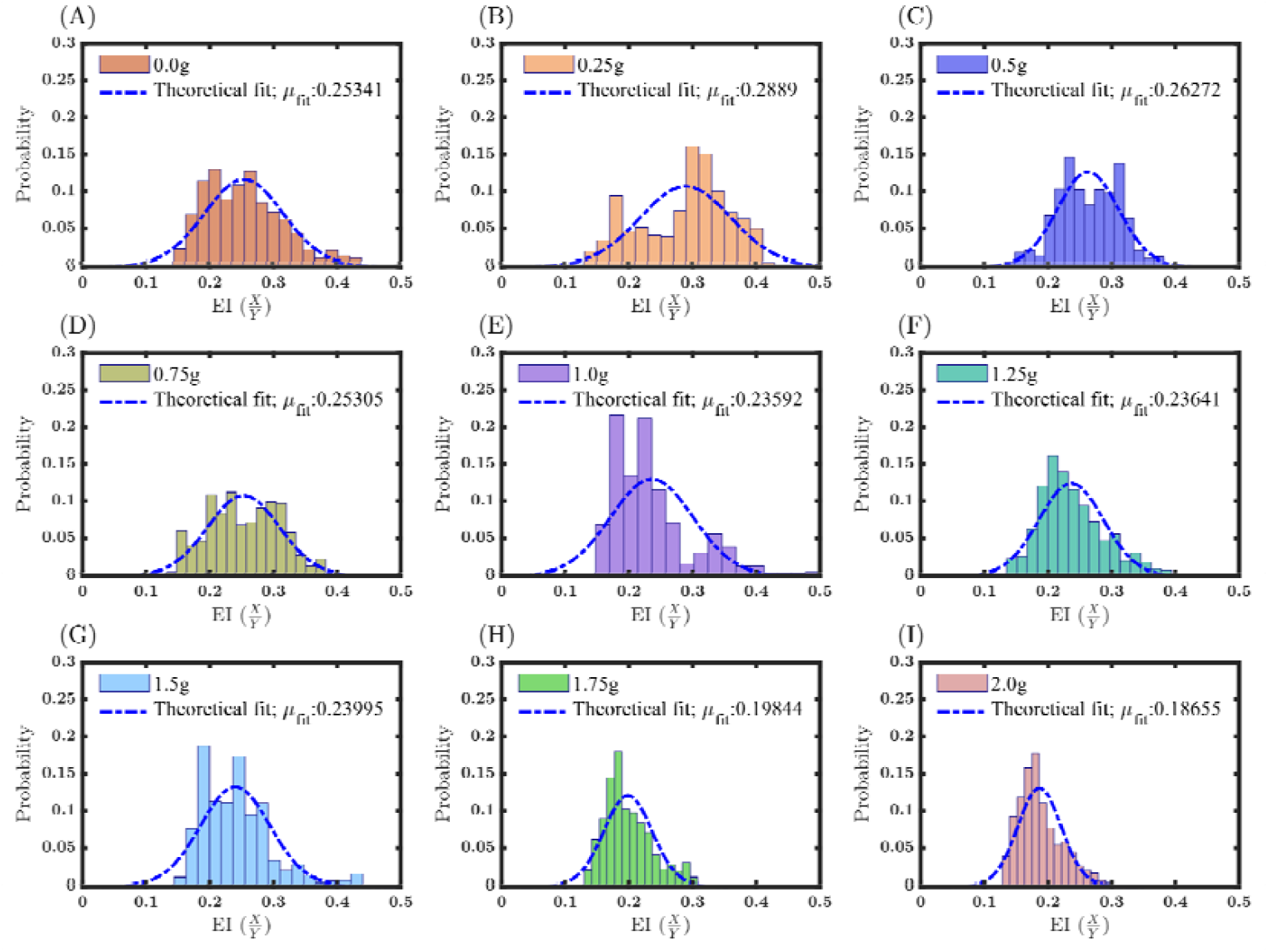
The probability distribution of the Elongation Index (EI) data. Fitted theoretical distribution over the EI data reveal a decreasing trend in the mean variation (A) 0.0g, (B) 0.25g, (C) 0.50g, (D) 0.75g, (E) 1.0g, (F) 1.25g, (G) 1.50g, (H) 1.75g and (I) 2.0g.

**Fig. 6:**
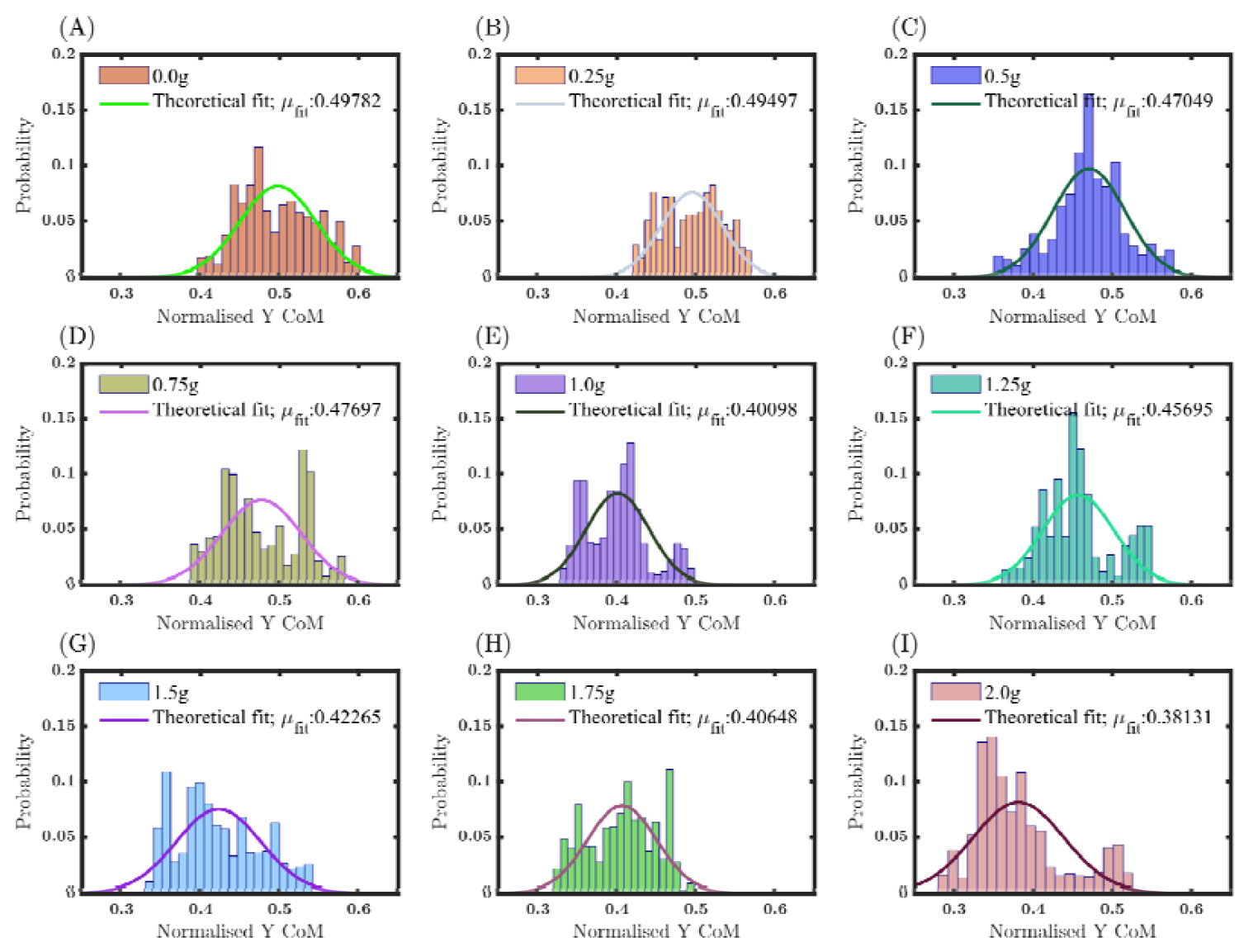
The probability distribution of the Y center of mass (YCoM) data. Fitted theoretical distribution over the YCoM data, comparing the mean of the distribution of 1g to the mean under other gravity conditions, the mean increases as the gravity decreases from 1g and decreases as the gravity condition increases (A) 0.0g, (B) 0.25g, (C) 0.50g, (D) 0.75g, (E) 1.0g, (F) 1.25g, (G) 1.50g, (H) 1.75g and (I) 2.0g.

The Mann-Whitney U test was conducted to validate the findings by aggregating 5112 samples for each gravitational group. We assumed the null hypothesis for the Mann-Whitney test to show no difference between the median of 1g and the tested gravitational condition. For the alternate hypothesis, the medians between the tested groups were supposed to be different. Significance levels were designated as p < 0.05, p < 0.01, and p < 0.001 to denote statistical significance between the compared samples.

**Fig. 7** compares the median values of EI, DI, Theta, and YCoM at 1g with those at other gravitational conditions. A noticeable difference in medians is observed between 0g and 2g conditions and substantial differences are evident in the central values across all gravitational conditions in the EI, DI, Theta and YCoM variables, as indicated by the confidence levels provided by the p-value. These findings support the alternate hypothesis, suggesting that gravitational variations induce changes in cell shape and facilitate spatial alignment in response to applied gravity.

**Fig. 7:**
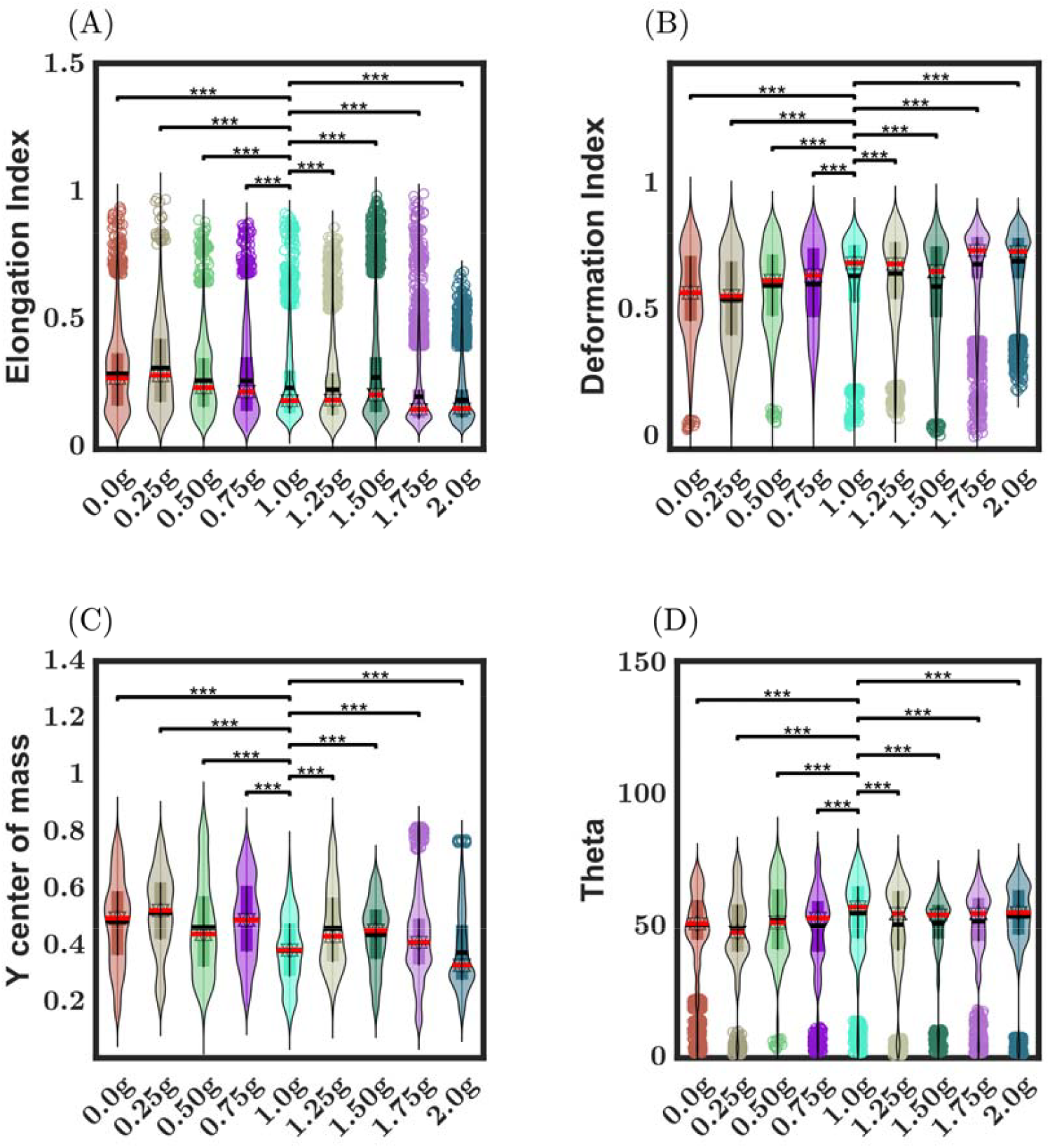
Violin plot illustrates the distribution of various solid parameters under different gravitational conditions (0.0g to 2.0g). The red bar represents the sample median, while the black bar indicates the mean. Statistical significance is denoted by asterisks: *p < 0.05, **p < 0.01, and ***p < 0.001. (A) Elongation Index (EI) demonstrates significant variations across gravity levels, reflecting changes in cell shape, with the null hypothesis (i.e., median at 1g equals sample median) being tested. (B) Deformation Index (DI) highlights notable differences in the degree of cellular deformation across conditions. (C) The Y Center of Mass (YCoM) exhibits a marked decline in median values as gravitational force increases. (D) Pitch angle (θ) shows a progressive increase with higher gravity levels.

In **Fig. 5**, while maintaining the mean of 1g distribution as a reference point, we assess the variation in the mean values. From the means of probability distribution plots, it is evident that the Elongation Index exhibits a linear relationship with the magnitude of gravitational force applied, starting from 0g to 2g. In order to provide additional verification, a linear regression model was employed. The coefficient of determination, denoted as R^2^, is calculated to be 0.8718 (**Fig. 8(A)**), suggesting that the model effectively describes the dataset. Moreover, the Pearson correlation coefficient was computed as −0.9337, suggesting a robust negative linear association between the observed data. A similar approach was employed for DI, theta, and normalized Y center of mass (YCoM). It is worth noting that the normalized Y center of mass exhibited a linear correlation with the magnitude of gravitational force applied, and their relationship was observed to be the same as that of the Elongation Index. The R^2^ for the fit was determined to be 0.7934 (**Fig. 8(B)**). The Pearson correlation coefficient was also calculated to be −0.8907, indicating a strong negative linear relationship between the observed data.

**Fig. 8:**
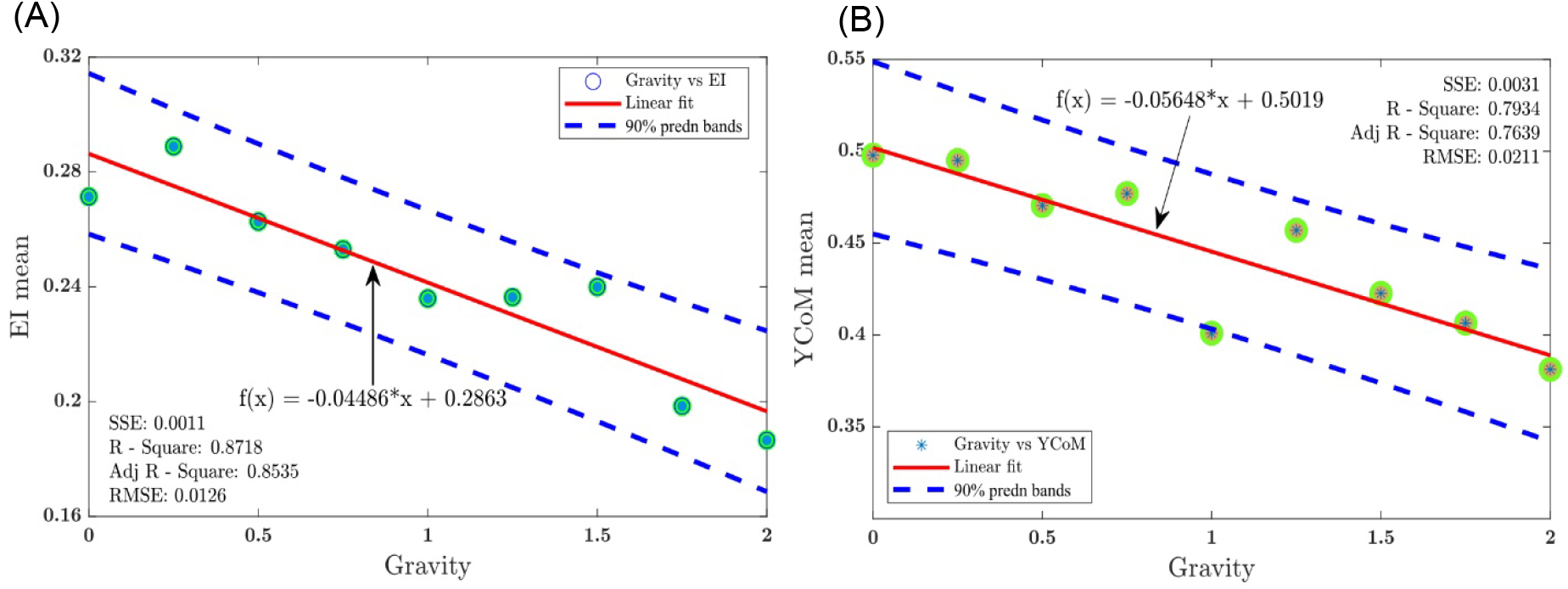
(A) Linear regression of mean distribution of EI with gravity. Fit shows a strong negative correlation between EI and applied gravity. (B) Linear regression of mean distribution of YCoM with gravity and fit shows a negative correlation between YCoM and applied gravity.

In addition to the temporal aggregation of solid forces (as shown in **Fig. 4**), parametric aggregation was performed; as illustrated in **Fig. 9**, a decrease in all three forces was observed during the hypergravity regime. In contrast, an increase was detected during the microgravity regime, with the most considerable increase occurring under the 0g condition. Upon reexamining **Eq. 22 to 25**, it becomes apparent that the magnitude of the drag force is dependent on the drag coefficient, which in turn is controlled by Reynold’s number. The Reynolds number is dependent on the magnitude of velocity. Velocity is a resultant quantity determined by the rate of change of an object’s acceleration, controlled by internal and external forces. The applied gravity is an external force acting in the y-direction with negative signage. It becomes clear that the external application of gravitational force becomes a subtractive force on the fluid and the solid, thereby decelerating it at higher g-levels.

**Fig. 9:**
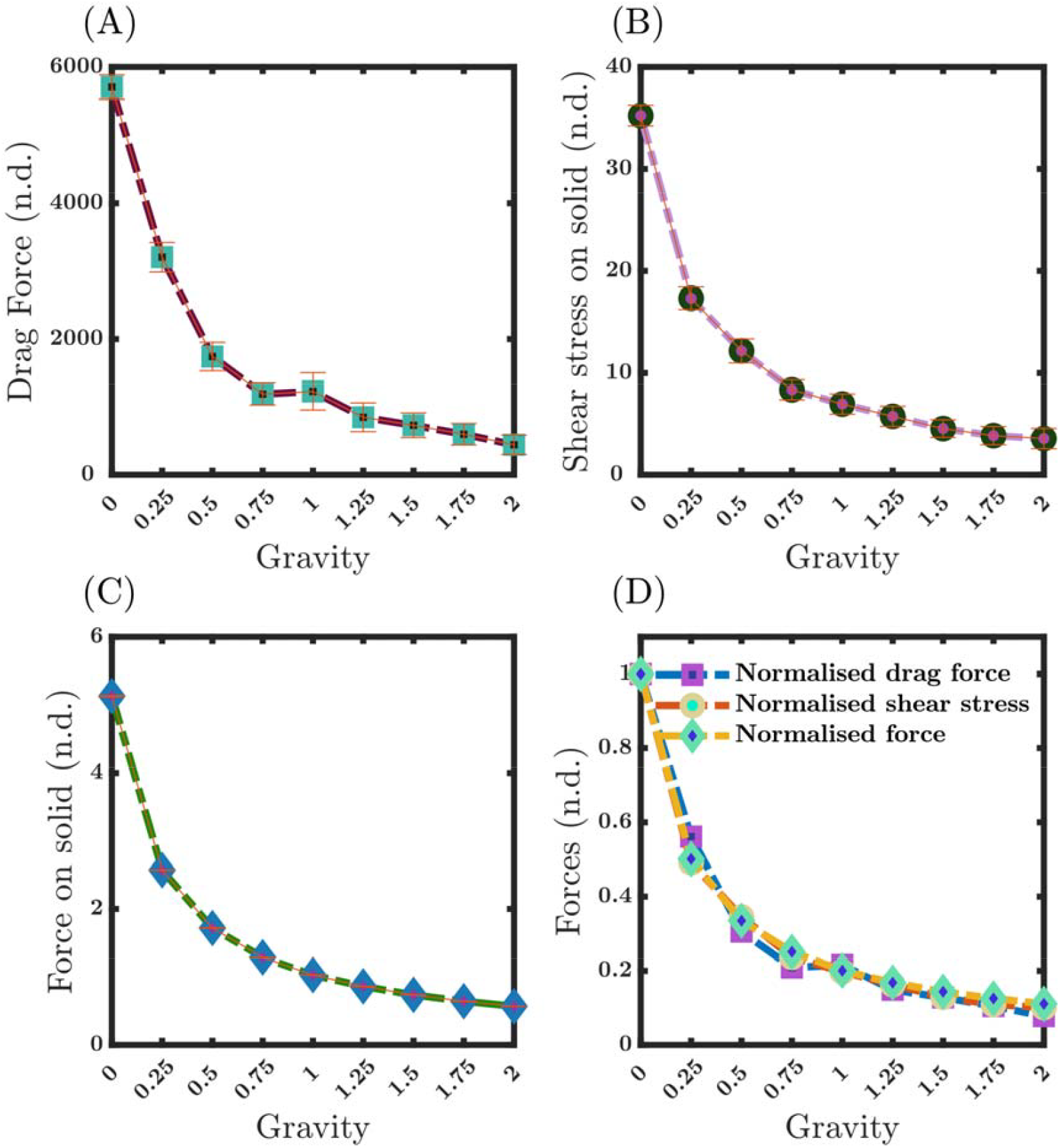
Parametric variation of forces acting on the Red Blood cell surface (n.d. denotes non-dimensionalized). (A) Drag forces shows marked decline with increasing gravity. (B) Shear stress on the solid surface follows a similar trend, decreasing with increasing gravity and stabilizing at higher gravity levels. (C) Force on solid declines with increasing gravity which is consistent with the trend of previous two forces. (D) Normalized forces, including drag force, shear stress, and overall force, exhibit consistent reduction patterns as gravity increases, stabilizing beyond 1.75g.

This study examined gravity’s impact on cells suspended in the bio-fluid medium. While previous research has explored the cellular interactions at the mesoscopic level using simulation (Fedosov et al. 2010; Xiao et al. 2014) and experimental approaches (Kumari et al. 2023; Tomaiuolo et al. 2011), the present study represents a novel investigation into the impact of gravitational conditions on cell morphology, distribution, and the corresponding forces. Previously, there have been some computational studies on the effects of microgravity on osteoclasts (Yang et al. 2018); however, there are no computational studies involving cells such as RBC and WBC in micro- or hypergravity regimes. Nevertheless, significant experimental advancements have been made on the impact of microgravity on vesicles and RBCs (Callens et al. 2008; Mader et al. 2006; Podgorski et al. 2011). To corroborate the results of our investigation, a comparative analysis was carried out between the detachment and lift of the RBC and vesicles under the 1g condition of Losserand et al. (Losserand et al. 2019) (refer to Figure 1(c)) with the present work. This study’s findings are consistent with the experimental data of the cell behavior near the wall (**Fig. 10: 1g**).

**Fig. 10:**
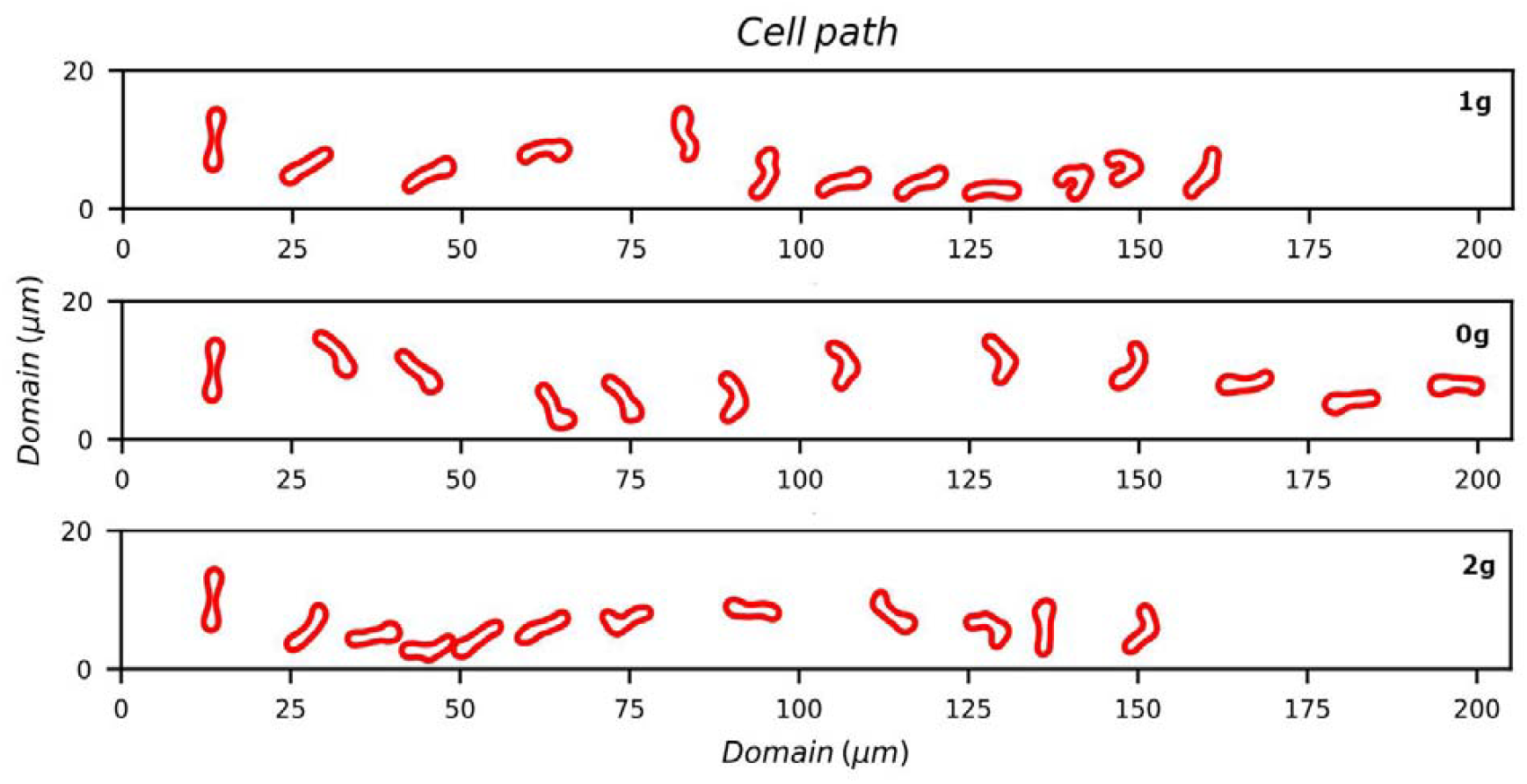
Distribution of the cell across the domain at selected g-conditions. Although the cell-free layer is formed at all three conditions, it is more pronounced under 0g with the cell pushed to the centerline due to higher lift force, whereas the cell under 1g and 2g moves closer to the wall.

As the cell approaches the wall, it experiences a lift force that results in its displacement away from the wall and towards the centerline. This results in a separation between the walls and the cell, commonly referred to as the cell-free layer. However, in the context of zero gravity, it was observed that the cell moved toward the wall even in the absence of gravitational forces and then pushed the cell with a greater upward force, enabling the formation of stable cell structures along the domain centerline (as depicted in (**Fig. 10: 0g**). In contrast, cells subjected to a gravitational force of 2g exhibit a significant proximity to the wall for longer durations, as depicted in **Fig. 10: 2g**. This phenomenon arises due to increased interaction with the nearby fluid particles, resulting in heightened viscous and dissipative forces. Consequently, the cell experiences a deceleration, thereby reducing drag, shear, and solid forces. Cell’s movement toward the domain wall also triggers the cells to change shape to accommodate the higher force near the wall.

## 4. Conclusion and Future Directions

This work has focused on the computational modeling of cell mechanics under an external gravitational force using Dissipative Particle Dynamics. Employing a discrete cellular structure model, the investigation revealed partial answers to initial inquiries regarding the cell’s response to changing gravitational forces. The findings reveal that RBC undergoes deformation and elongation in response to gravitational variations, with noticeable alterations in measures such as the EI and DI compared to the 1g condition (as indicated by a p-value < 0.001) signifying shape alteration and spatial realignment. At the same time, the normalized center of mass on the y-axis decreased, indicating a downward shift within the blood vessel. Decreasing normalized center of mass raises the question of how different forces are concentrated on the cell. Solid forces, including drag and shear stress, were scrutinized to answer that. Solid forces include the forces that are exerted on the solid by the fluid within a neighborhood. These forces experience a reduction in their magnitude whenever there is an increase in the gravity of the cell, mainly a reduction of the cell velocity due to its non-adherent nature. Notably, the study illuminated the dynamic morphological transformations of red blood cells (RBCs) from a biconcave shape to various configurations shaped by their spatial orientation and interaction with the surrounding fluid. The findings of this study have important implications for understanding cell mechanics and the physical response of cells to external forces.

Through numerical simulations, this study pioneers exploring microgravity’s influence on cells suspended in a bio-fluid. Future endeavors will involve scaling the research to encompass physiologically relevant quantities of RBCs and immune cells and subject them to varying gravity. Thereby offering crucial insights into fundamental cellular behaviors. Further investigations into the specific molecular and biomechanical mechanisms underlying these observed responses will be pivotal for advancing our comprehension of cellular adaptations to varying gravitational environments.

## 5 Acknowledgements

RRS acknowledges funding from the Department of Biotechnology, Govt. of India (Sanction Order No. BT/PR35683/BID/7/937/2019, dated 17.12.2020); AM acknowledges CSIR-GATE for Senior Research Fellowship. The support and the resources provided by PARAM Brahma Facility under the National Supercomputing Mission, Government of India at the Indian Institute of Science Education and Research; Pune are gratefully acknowledged.

## 7. Author contributions

AM performed coding, simulation, analysis and writing; RRS conceived the study, critical analysis, writing, proofreading and acquired funding.

## 8. Data and code availability

Codes and datasets generated during the current study will be available upon request.

## 10. Ethics, Consent to Participate, and Consent to Publish declarations

Not applicable.

