## Supplementary Information for "Impact of gravitational forces on Red Blood Cell dynamics in biofluid suspension"

### Section 1 - Parameter derivation:

To ensure coherence between the simulation system and the actual system, it is essential to establish a correspondence between the physical attributes and the dimensionless features within the model. The scaling procedure serves to establish a relationship between the nondimensional units utilized in the model and their equivalent physical units. The scales are provided in the following manner:

Length scale = $\frac{L^{P}}{L^{M}}$ [m]

Energy scale = $\frac{\boldsymbol{Y}^{\boldsymbol{P}}}{\boldsymbol{Y}^{\boldsymbol{M}}}\left( \frac{\boldsymbol{L}^{\boldsymbol{P}}}{\boldsymbol{L}^{\boldsymbol{M}}} \right)^{\boldsymbol{2}}$[Nm]

Force scale = $\frac{\boldsymbol{Y}^{\boldsymbol{P}}}{\boldsymbol{Y}^{\boldsymbol{M}}}\left( \frac{\boldsymbol{L}^{\boldsymbol{P}}}{\boldsymbol{L}^{\boldsymbol{M}}} \right)$ [N]

Body force scale = $\frac{Y^{P}}{Y^{M}}\left( \frac{L^{M}}{L^{P}} \right)^{2}(\frac{\rho_{DPD}}{\rho_{Plasma}})$ [m/s^2^]

Time scale = $\frac{L^{P}}{V^{P}}$ [s]; superscripts M and P denote “model” and “physical”.

### Section 2 - Figures:

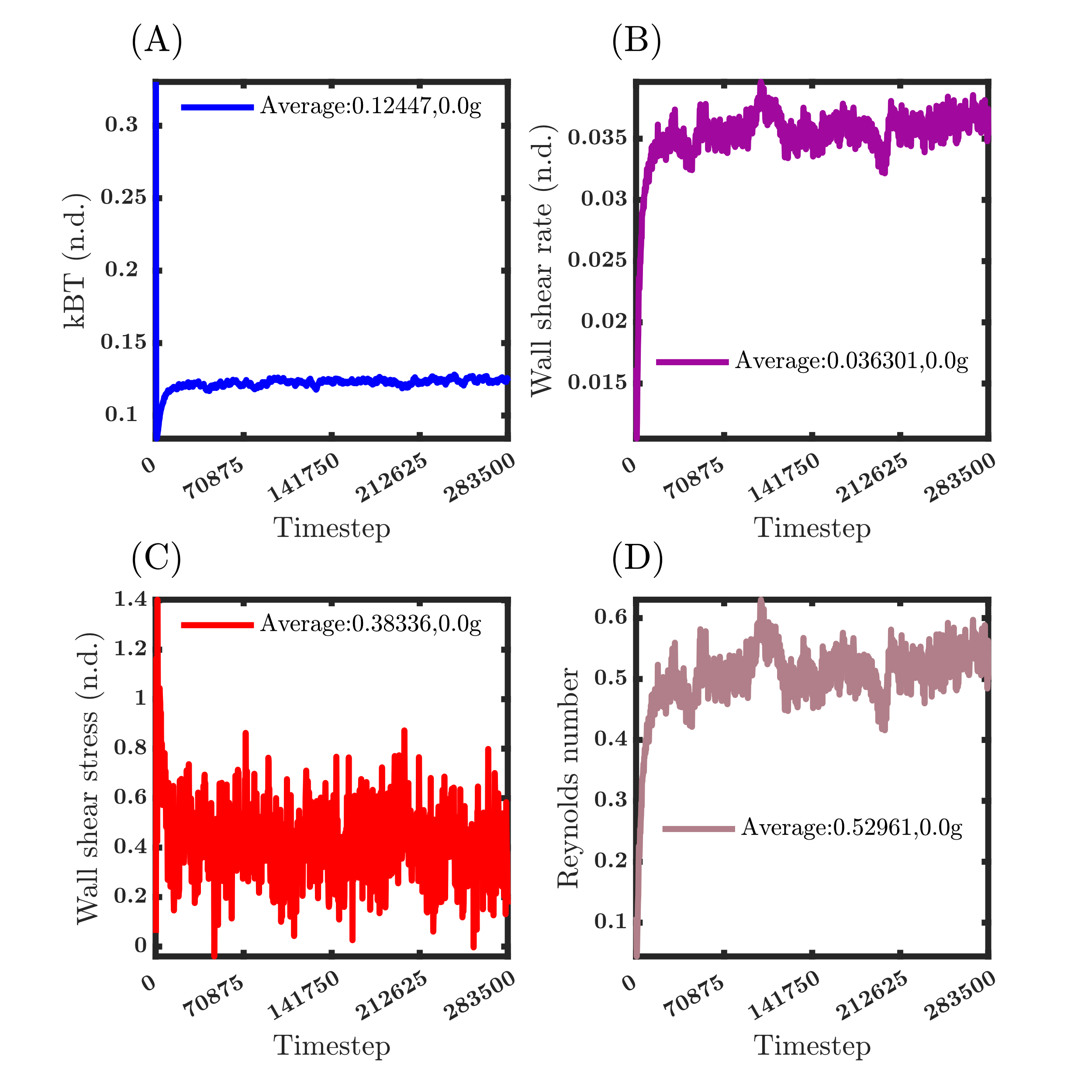

**Fig. S1: Temporal evolution of various parameters during the simulation under 0g condition; value averaged for the last 25% of data. (A) Temperature fluctuations (kBT) stabilize after 10,000 timesteps. (B) The wall shear rate fluctuates eventually stabilizes. (C) Wall shear stress rises at the start of the simulation and eventually reaches equilibrium. (D) Reynolds number initially increases and eventually stabilizes.**

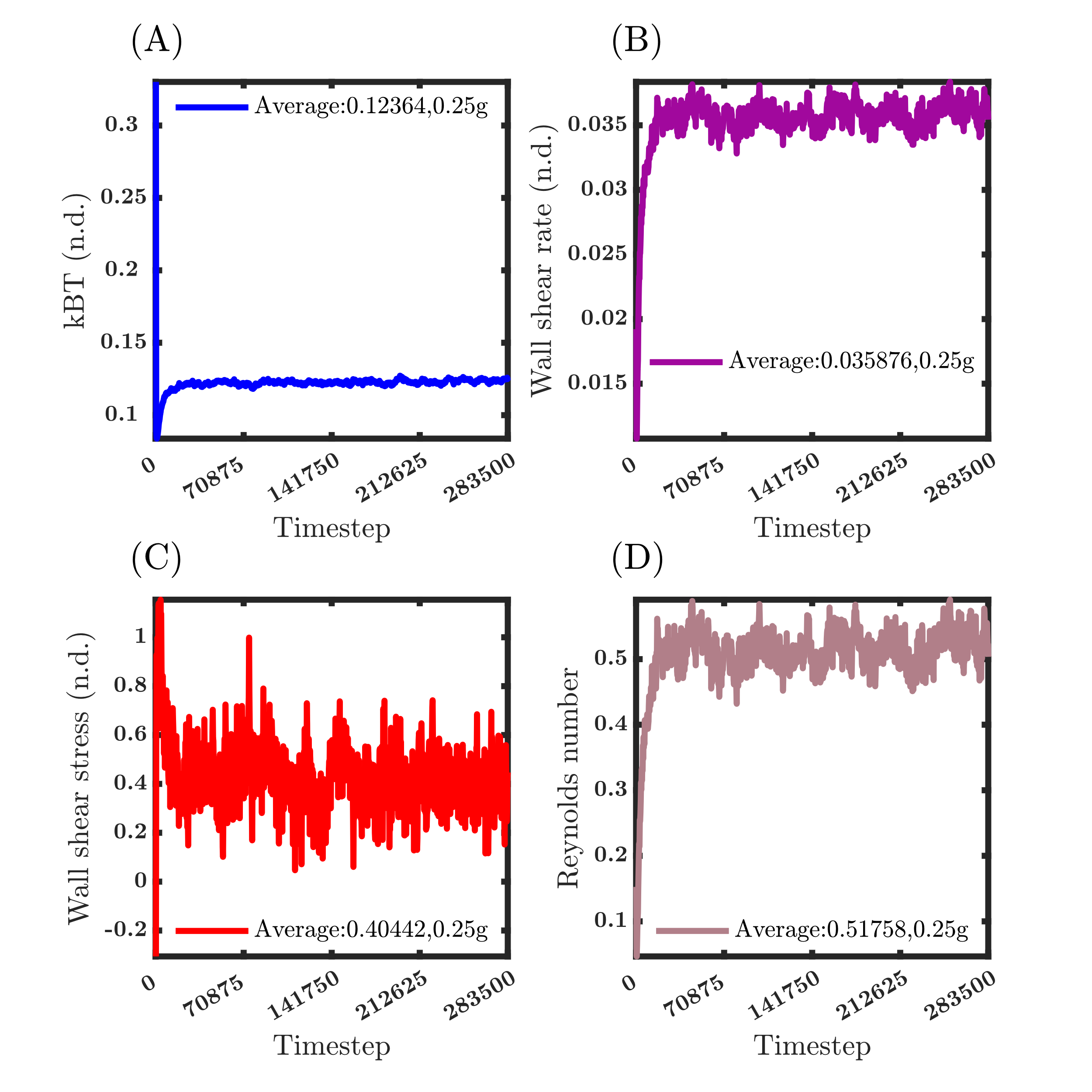

**Fig. S2: Temporal evolution of various parameters during the simulation under 0.25g condition; value averaged for the last 25% of data. (A) Temperature fluctuations (kBT) stabilize after 10,000 timesteps. (B) The wall shear rate fluctuates eventually stabilizes. (C) Wall shear stress rises at the start of the simulation and eventually reaches equilibrium. (D) Reynolds number initially increases and eventually stabilizes.**

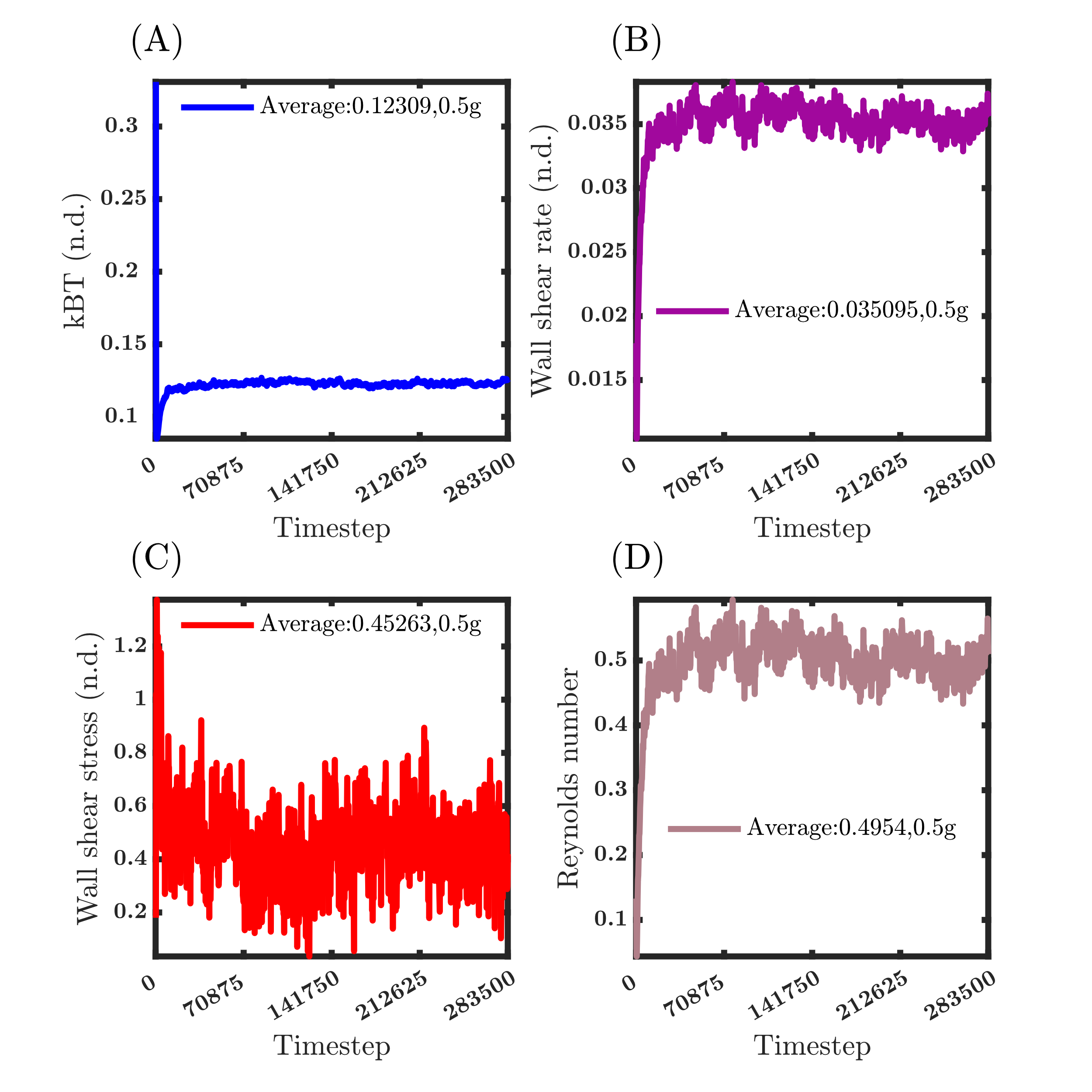

**Fig. S3: Temporal evolution of various parameters during the simulation under 0.5g condition; value averaged for the last 25% of data. (A) Temperature fluctuations (kBT) stabilize after 10,000 timesteps. (B) The wall shear rate fluctuates eventually stabilizes. (C) Wall shear stress rises at the start of the simulation and eventually reaches equilibrium. (D) Reynolds number initially increases and eventually stabilizes.**

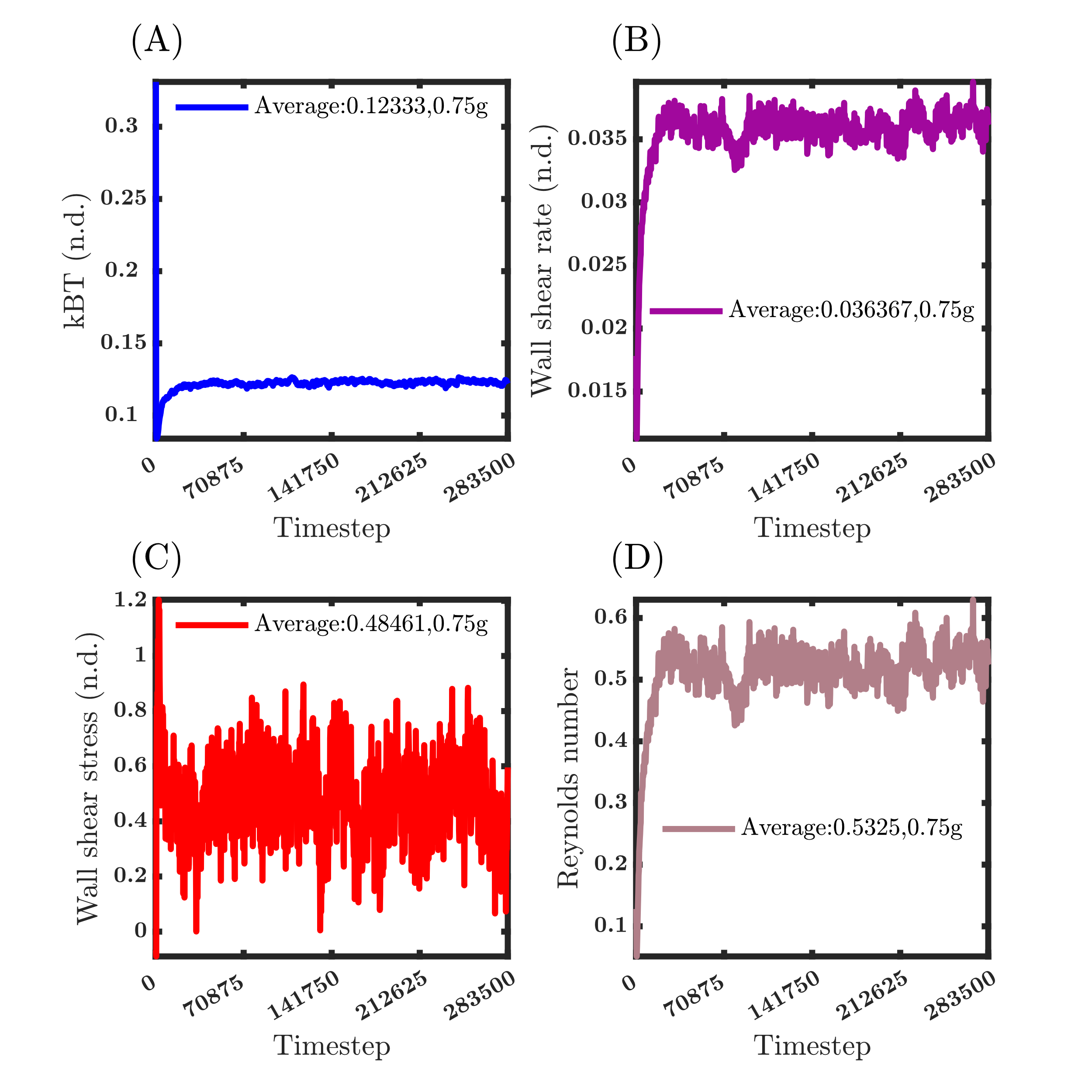

**Fig. S4: Temporal evolution of various parameters during the simulation under 0.75g condition; value averaged for the last 25% of data. (A) Temperature fluctuations (kBT) stabilize after 10,000 timesteps. (B) The wall shear rate fluctuates eventually stabilizes. (C) Wall shear stress rises at the start of the simulation and eventually reaches equilibrium. (D) Reynolds number initially increases and eventually stabilizes.**

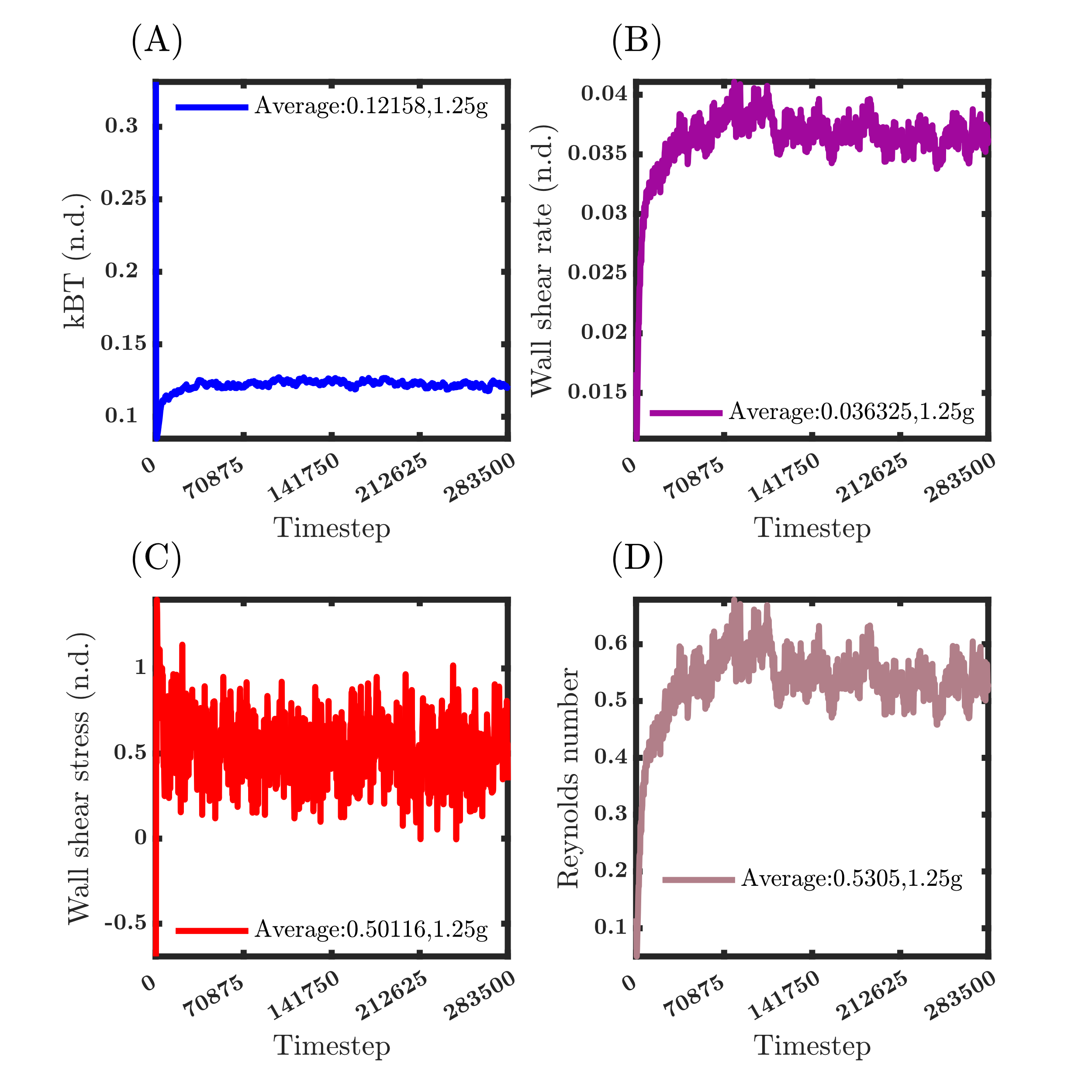

**Fig. S5: Temporal evolution of various parameters during the simulation under 1.25g condition; value averaged for the last 25% of data. (A) Temperature fluctuations (kBT) stabilize after 10,000 timesteps. (B) The wall shear rate fluctuates eventually stabilizes. (C) Wall shear stress rises at the start of the simulation and eventually reaches equilibrium. (D) Reynolds number initially increases and eventually stabilizes.**

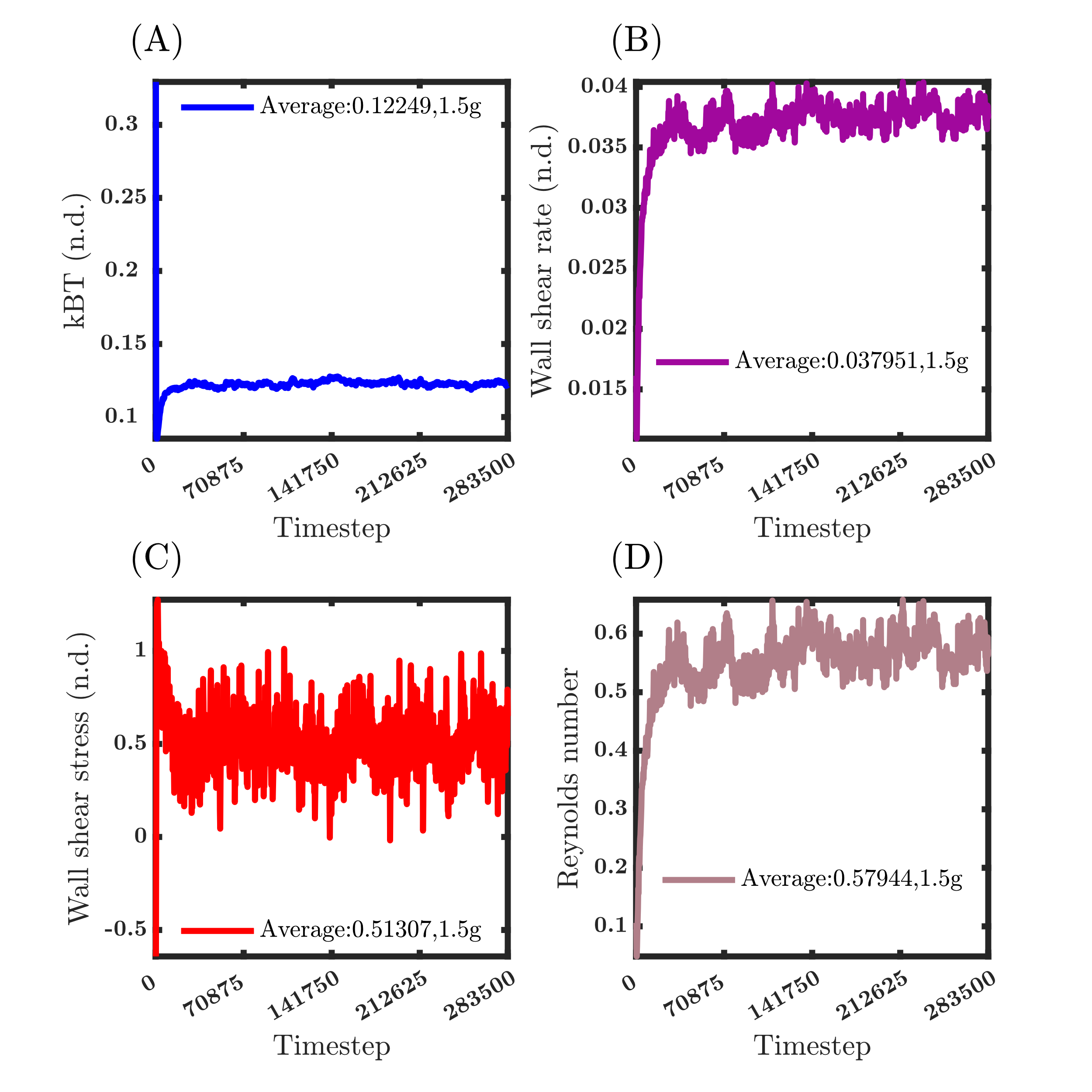

**Fig. S6: Temporal evolution of various parameters during the simulation under 1. 50g condition; value averaged for the last 25% of data. (A) Temperature fluctuations (kBT) stabilize after 10,000 timesteps. (B) The wall shear rate fluctuates eventually stabilizes. (C) Wall shear stress rises at the start of the simulation and eventually reaches equilibrium. (D) Reynolds number initially increases and eventually stabilizes.**

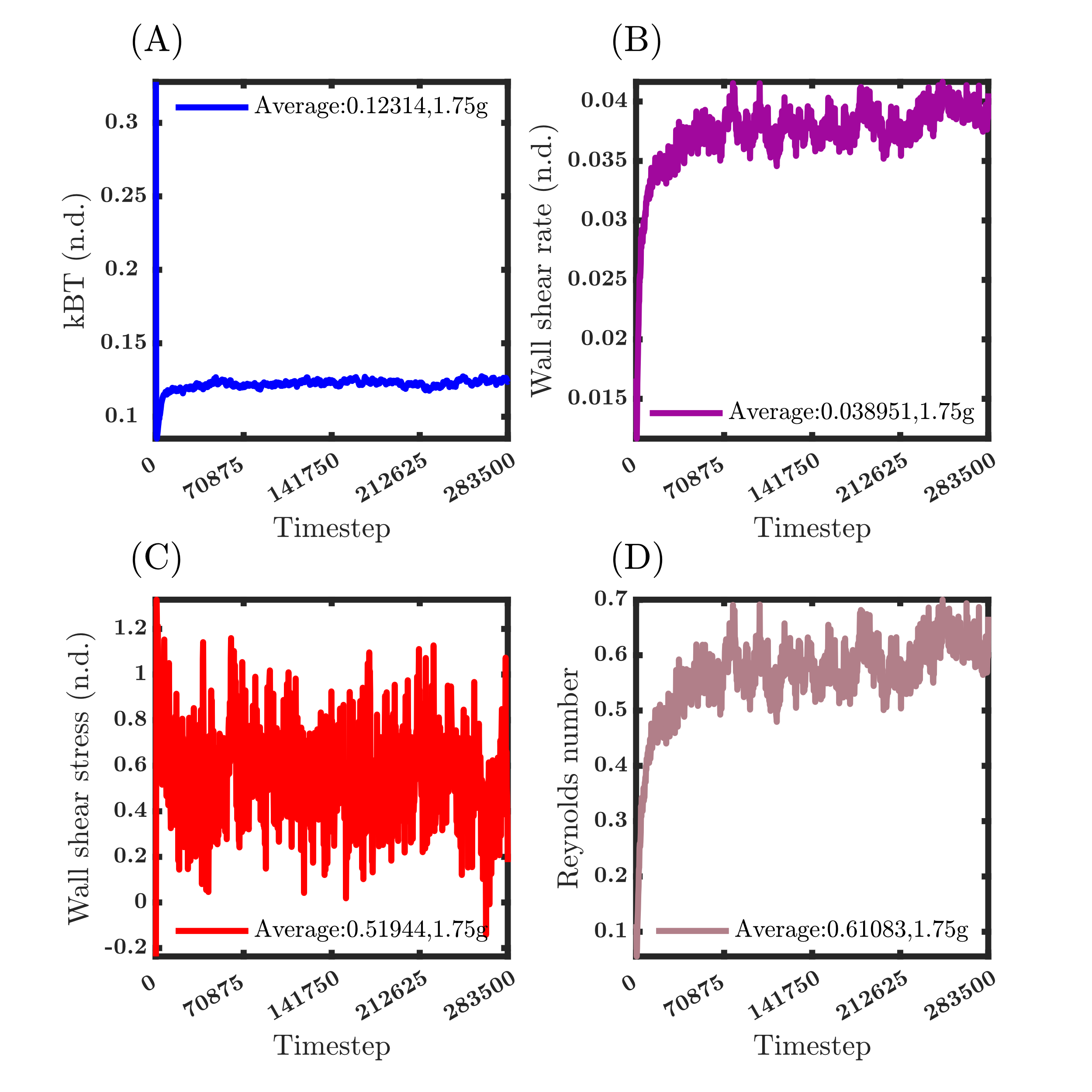

**Fig. S7: Temporal evolution of various parameters during the simulation under 1.75g condition; value averaged for the last 25% of data. (A) Temperature fluctuations (kBT) stabilize after 10,000 timesteps. (B) The wall shear rate fluctuates eventually stabilizes. (C) Wall shear stress rises at the start of the simulation and eventually reaches equilibrium. (D) Reynolds number initially increases and eventually stabilizes.**

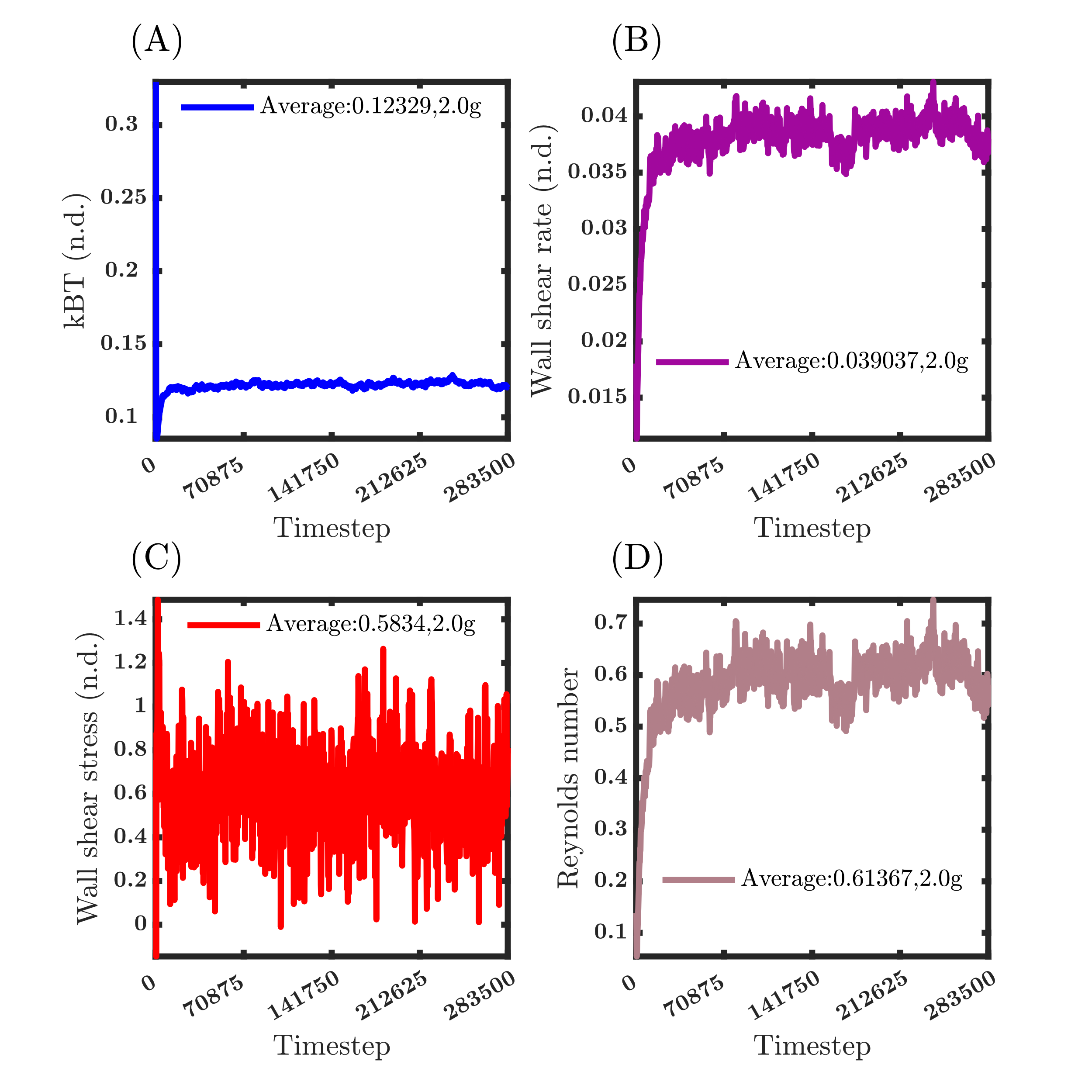

**Fig. S8: Temporal evolution of various parameters during the simulation under 2.00g condition; value averaged for the last 25% of data. (A) Temperature fluctuations (kBT) stabilize after 10,000 timesteps. (B) The wall shear rate fluctuates eventually stabilizes. (C) Wall shear stress rises at the start of the simulation and eventually reaches equilibrium. (D) Reynolds number initially increases and eventually stabilizes.**

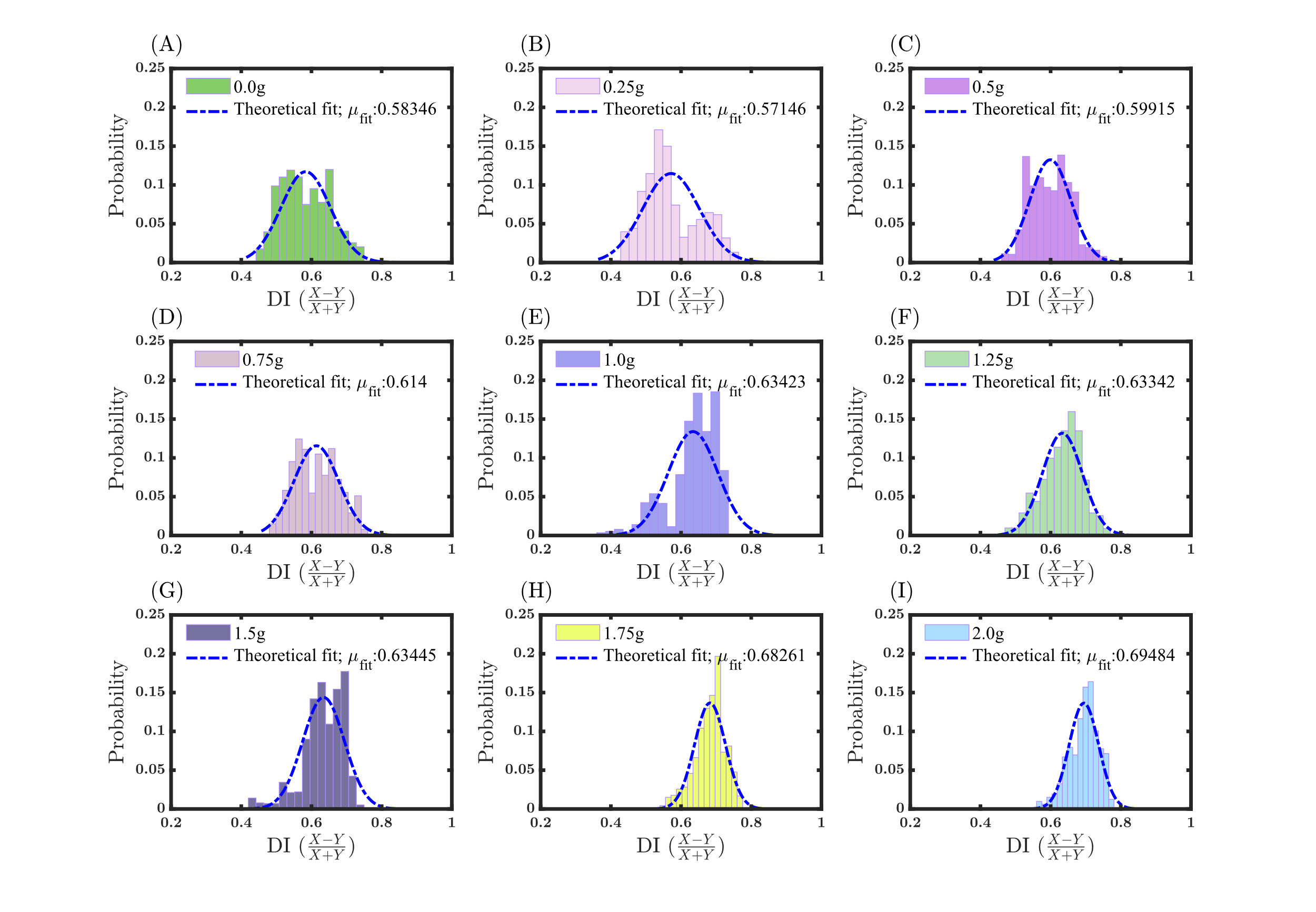

**Fig. S9: The probability distribution of the Deformation Index (DI) data. Fitted theoretical distribution over the DI data reveal a clear trend of increasing deformation under hypergravity and decreasing deformation under microgravity when compared with 1g. (A) 0.0g, (B) 0.25g, (C) 0.50g, (D) 0.75g, (E) 1.0g, (F) 1.25g, (G) 1.50g, (H) 1.75g and (I) 2.0g.**

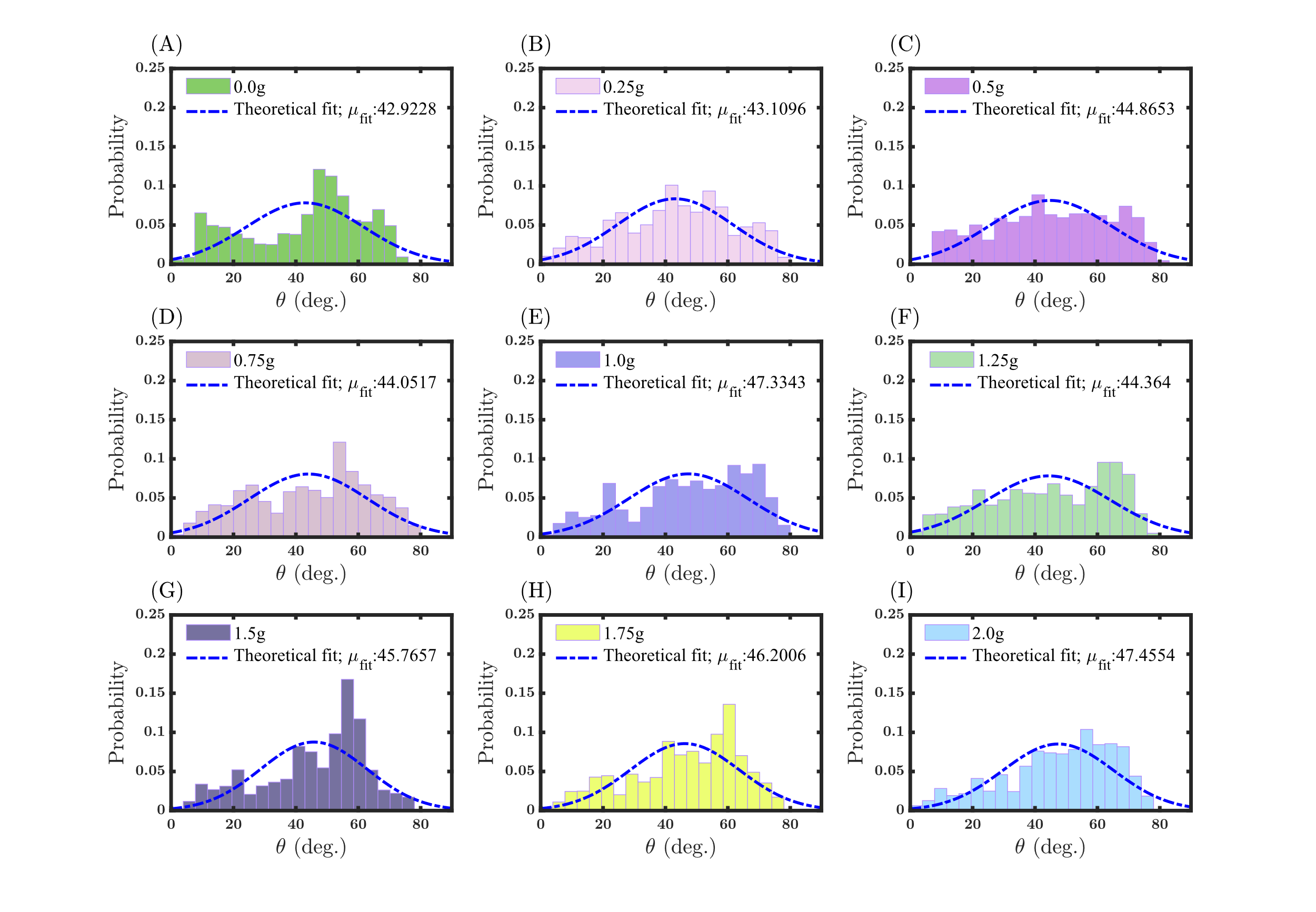

**Fig. S10: The probability distribution of the Pitch angle (Theta). Fitted theoretical distribution over the theta data reveal an increasing trend of theta to the applied gravity (A) 0.0g, (B) 0.25g, (C) 0.50g, (D) 0.75g, (E) 1.0g, (F) 1.25g, (G) 1.50g, (H) 1.75g and (I) 2.0g.**
